# Transcriptomic disruption and functional hypoactivity in DYT-*SGCE* MGE-patterned inhibitory neurons

**DOI:** 10.1101/2024.12.19.629430

**Authors:** Zongze Li, Laura Abram, Maria Cruz-Santos, Olena Petter, Kathryn J Peall

## Abstract

Myoclonus Dystonia is a dystonic movement disorder caused by *SGCE* mutations, the underlying pathophysiology for which remains unclear. Here, we evaluated the impact of *SGCE* mutations on medial ganglionic eminence (MGE)-derived GABAergic neurons using patient-derived induced pluripotent and gene edited embryonic stem cell lines, each compared to their isogenic wild-type control. No significant differences were observed in markers of neuronal development however, single-cell RNA sequencing demonstrated transcriptomic dysregulation in genes related to axonal organization, synaptic signalling, and action potential generation in the *SGCE* -mutation harbouring neurons. Functional assays demonstrated reduced neurite outgrowth, lower calcium responses to GABA, and decreased neuronal excitability and network activity in the *SGCE* -mutant neurons. These findings contrast with the hyperexcitable phenotype previously observed in *SGCE* -mutant cortical glutamatergic neurons. Collectively, this supports loss of neuronal inhibitory activity, and disruption to the neuronal excitatory/inhibitory balance in motor circuits, in contributing to the overall hyperkinetic clinical phenotype in Myoclonus Dystonia.

## Introduction

Dystonia is one of the most common forms of movement disorder, with an estimated population prevalence of 1.2%.^1,2^ It involves loss of co-ordinated contraction of antagonistic muscle groups, leading to abnormal postures and pain, with subsequent impact on quality of life.^3^ The clinical presentation of dystonia is heterogenous, involving single or multiple muscle groups (focal, segmental or generalized), genetic or idiopathic in aetiology, and of childhood or adult onset.^4,5^ Co-morbid psychiatric symptoms are also observed, frequently pre-dating onset of the motor symptoms and spanning a spectrum of anxiety, obsessive-compulsive disorder (OCD), depression and social phobia.^6^

In excess of 50 Mendelian-inherited dystonia-causing genes have now been identified, with these predominantly resulting in onset of motor symptoms in early life stages.^7^ One such disorder is Myoclonus Dystonia, caused by mutations in the autosomal dominantly inherited *SGCE* gene, encoding the ε-sarcoglycan protein.^8,9^ The associated clinical phenotype typically involves upper body predominant myoclonus, focal or segmental dystonia involving the cervical and/or upper limb regions, and psychiatric symptoms including generalised anxiety disorder and OCD.^10,11^ The ε-sarcoglycan protein is a single-pass transmembrane glycoprotein,^12^ which is expressed embryonically and postnatally, suggesting its importance in development.^13^ Examination of the brain-specific form of ε-sarcoglycan suggests it forms part of a brain-specific dystrophin-associated protein complex,^14^ with ultra-deep sequencing of post-mortem brain tissue demonstrating high levels of expression in the primary somatosensory cortex.^15^

Understanding of the pathophysiological mechanisms underpinning dystonia remain limited however, evidence from human imaging, murine and post-mortem studies indicate a disruption to neuronal networks, principally involving the basal ganglia-cerebello-thalamo-cortical circuits.^16,17^ Human imaging studies have identified white and grey matter morphometric and white matter microstructural changes in regions linked with the cerebral cortex,^18–21^ and electrophysiological studies, in humans and mice, have demonstrated increased cortico-striatal long-term potentiation (LTP) and reduced long-term depression (LTD) in the motor cortex.^22–24^ Several studies have suggested that impaired cortical surround inhibition is responsible for these changes however, recent work has suggested a more complex picture with additional factors contributing to the hyperexcitable phenotype.^24,25^ Our recent work involving CRISPR-edited embryonic stem cell and patient derived induced pluripotent stem cell (iPSC) models of *SGCE* -mutation positive Myoclonus Dystonia differentiated towards an excitatory glutamatergic cortical neuronal lineage, demonstrated a hyperexcitable phenotype, more complex dendritic branching morphology and disruption to synaptic adhesion molecules neurexin-1 and neuroligin-4.^2^

This study seeks to build on this work through differentiation of the same stem cells models towards an MGE-like-derived inhibitory neuronal lineage, the most common subtype of inhibitory neurons in the cerebral cortex, determining the impact of the loss of ε-sarcoglycan expression on their development, structure and function. We aim to determine whether the excess motor activity observed in dystonia is driven purely by the hyperexcitability observed in excitatory glutamatergic cortical neurons, or whether, in line with *in vivo* electrophysiological evidence, there is an additional loss of inhibitory activity in MGE-derived GABAergic neurons. Understanding these changes are critical to identification of novel therapeutic targets and future therapeutic development.

## Results

### Generation of MGE-derived GABAergic neurons

To investigate the role of MGE-derived GABAergic neurons in Myoclonus Dystonia, neurons were differentiated from two patient-derived *SGCE* mutation-positive hiPSC lines (Patient 1 and 2) and a single CRISPR/Cas9 edited *SGCE* knock-out-iCas9 human embryonic stem cell (hESC) and their matched isogenic wild-type control lines (Figure 1A). Immunocytochemical staining at day 25 of differentiation demonstrated no significant differences in the proportion of cell expressing key MGE progenitor markers, NKX2.1 and FOXG1, anterior forebrain marker, OTX2, and neural stem cell marker, NESTIN between each cell line pair (Figure 1B-C). At the same time point, markers of ventral midbrain (FOXA2), caudal ganglionic eminence (NR2F2), dorsal forebrain and lateral ganglionic eminence (PAX6) were detected in a minimal proportion of cells with no significant differences between lines (Figure 1c, Supplementary Figure 1A). At day 80 differentiation, comparable levels of neuronal (NEUN and TUJ1), GABAergic (GAD67 and GABA), and MGE lineage (FOXG1, OLIG2, and SOX6) markers were observed across all cell lines, again with no significant difference between mutant and wild-type lines for each cell line pair (Figures 1D-I). Some neurons co-expressed SST and CB, suggesting the presence of GABAergic neuron subtypes (Figure 1J). Further examination of gene expression using qPCR similarly revealed an upregulation of MGE-derived GABAergic neuronal markers (*LHX6*, *GAD67*, *NKX2.1* and *SST*) at day 80 of the differentiation protocol (Supplementary Figure 1B).

**Figure 1.**
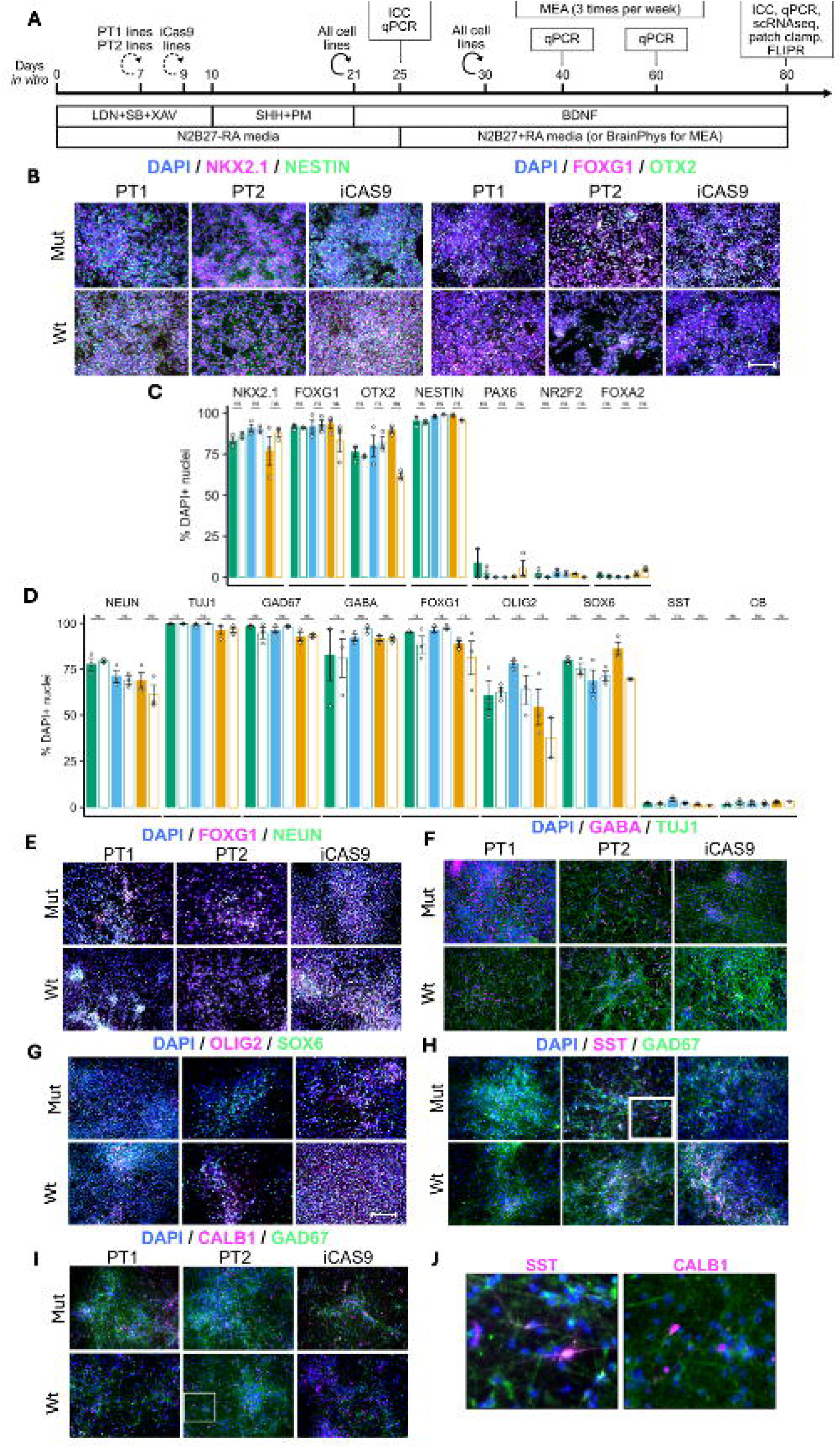
Absence of impact of *SGCE* mutations on differentiation and developmental markers of medial ganglionic eminence-derived GABAergic neurons. **A:** Schematic overview of the differentiation protocol used to derive medial ganglionic eminence (MGE)-GABAergic neurons and study design. **B:** Day 25 differentiation: representative immunofluorescence images for MGE progenitor markers (NKX2.1, FOXG1), anterior forebrain marker (OTX2), neural stem cell marker (NESTIN). Scale bar[=[100[µm. **C:** Day 25 differentiation quantification of immunofluorescent markers NKX2.1, FOXG1, OTX2, NESTIN, and non-MGE lineage markers: PAX6, NR2F2, and FOXA2. **D:** Day 80 differentiation quantification of immunofluorescent markers of neuronal markers (NEUN, TUJI), GABAergic neuronal markers (GAD67, GABA), markers of MGE-lineage (FOXG1, OLIG2, SOX6) and GABAergic neuronal subtypes (SST, CB). Day 80 differentiation representative immunofluorescence images for **E:** FOXG1 and NEUN, **F:** GABA and TUJ1, **G:** OLIG2 and SOX6, **H:** SST and GAD67, **I:** CALB1 and GAD67. **J:** Zoomed views of SST and CALB1 immunofluorescence markers for the two regions highlighted by the white boxes. Scale bar[=[100[µm. Data presented as mean±SEM from three independent experiments. Lines compared using Wilcoxon signed-rank tests with FDR correction. ns: not statistically significant (p>0.05), *p<0.05. **p<0.01, ***p<0.001. BDNF: brain derived neurotrophic factor; ESC: embryonic stem cell; ICC: immunocytochemistry; iPSC: induced pluripotent stem cell; LDN: LDN193189; MEA: multi-electrode array; MGE: medial ganglionic eminence; NPC: neural progenitor; PM: purmorphamine; PT1: patient #1; PT2: patient #2; RGL: radial glia; SB: SB431542; scRNAseq: single-cell RNA-sequencing; SHH: Sonic Hedgehog; XAV: XAV939

Data captured from the single-cell RNA-sequencing (scRNAseq) analysis was used for further confirmation of cell line genotype and demonstration of markers consistent with MGE-derived GABAergic neurons. Here, *SGCE* mutation-carrying lines demonstrated significantly lower *SGCE* expression while no significant difference was observed in the expression levels of the other members of the sarcoglycan family (Supplementary Figure 1C). Following data integration (Supplementary Figure 2A), seven clusters of conserved cell types were identified across the two genotypic groups (Figure 2A, Supplementary Figure 2B). Two clusters (Neuron1 and Neuron2) expressed high levels of pan-neuronal markers (*SYT1*, *STMN2*, *SNAP25*, *RBFOX3*, and *DCX*) but low levels of progenitor (*MKI67*, *TOP2A*, *NES*, *SOX1*), radial glia (*SLC1A3*, *RFX4*, *FABP7*), and glial (*SOX10*, *PDGRFA*, *GJA1*, *GFAP*, *CD44*, *AQP4*) markers. In addition, Neuron1 and Neuron2 clusters expressed higher levels of markers for cells of an MGE-derived lineage (*NKX2-1*, *LHX6*, *FOXG1*, *MAFB*, *ERBB4*, *ARX*, *SOX6*) and GABAergic neurons (*GAD1*, *SLC32A1*, *CALB1*, *RELN*, *SST*, *NPY*). There was a near absence of markers for cells of lateral and caudal ganglionic eminence lineage, as well as glutamatergic, dopaminergic and cholinergic neurons (Figure 2B). In addition, SCENIC regulon enrichment found Neuron1 and Neuron2 clusters to be differentially enriched for regulons of key transcription factors involved in the specification of MGE-derived GABAergic neurons (including LHX6, SOX6, NKX2.1, FOXG1, ARX, MAFB, MEF2C, DLX5/6) but not transcription factors involved in the development of other neuronal lineages (Figure 2C).

**Figure 2:**
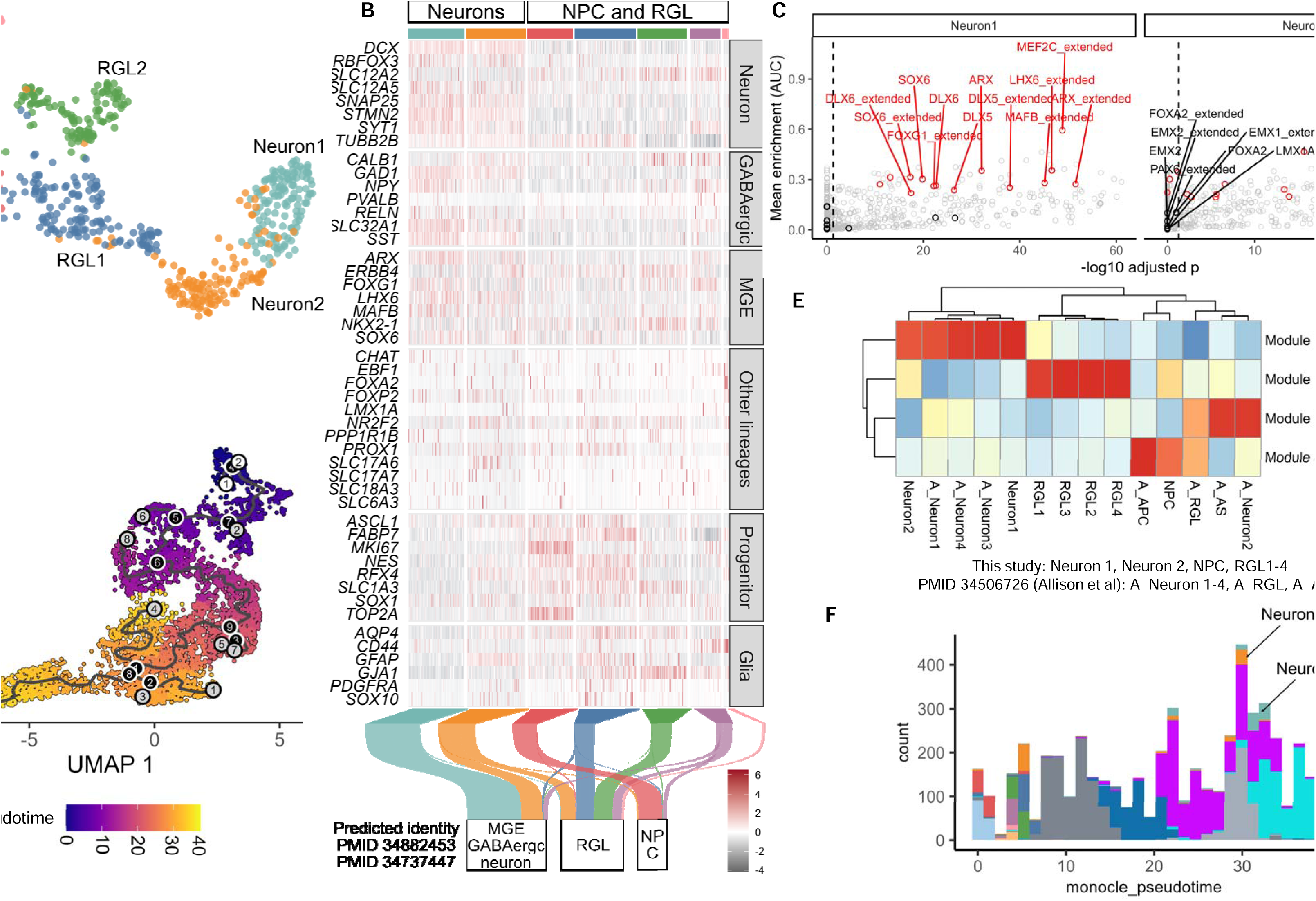
Transcriptomic support for the derivation of MGE-derived GABAergic neurons with maturation patterns consistent with *in vivo* derived data. **A:** D80 Uniform Manifold Approximation and Projection (UMAP) plot of single-cell RNA-sequencing (scRNAseq) of day 80 wild-type and *SGCE* - mutation positive MGE-derived GABAergic neurons coloured by cluster annotations. **B:** Heatmap of the expression of canonical marker genes across distinct lineages and cell types, present by cluster. Sankey plot indicating the predicted identity of each cell based on two previously published human foetal MGE scRNAseq datasets. **C:** Differentially enriched regulons in neuronal clusters highlighting regulons related to MGE development (red) and non-MGE (black). **D:** UMAP plot delineating the pseudo-temporal trajectory. Colour gradient references indicate the pseudotime score. **E:** Averaged expression of gene modules dynamically regulated along the pseudo-temporal trajectory grouped by cell-type clusters identified in this study and previously reported data. **F:** Histogram representing the pseudotime scores of cells across the integrated PSC-derived dataset, with Neuron 1 and Neuron 2 clusters identified in this study labelled. Key: APC: astrocyte progenitor cell; MGE: medial ganglionic eminence; NPC: neural progenitor; RGL: radial glia; UMAP: uniform manifold approximation and projection

Two publicly available scRNAseq datasets of human foetal ganglionic eminence tissue were used as reference to ensure that the GABAergic neurons generated resembled those found *in vivo*. Here, cells in clusters Neuron1 and Neuron2 were predicted to resemble MGE-derived GABAergic neurons in both datasets, while NPC clusters and the four radial glial clusters mapped to neural progenitors, radial glial and glial populations, respectively (Figure 1B, Supplementary Figure 2C). Moreover, integrative analysis revealed that the neurons generated exhibited similar human foetal MGE-derived GABAergic neuron-like transcriptomic identity to PSC-derived MGE-lineage GABAergic neurons observed in previously reported data (Supplementary Figure 3A-C).^25^ Furthermore, to compare the transcriptomic maturity between PSC-derived MGE-lineage GABAergic neurons generated and those reported in Allison et al., a pseudotemporal trajectory was constructed (Figure 2D). Genes upregulated along the trajectory (module 3) included genes related to aspects of neuronal development and maturation (Figure 2E, Supplementary Figure 3D, Supplementary Data 1), suggesting that the trajectory represents a neurodevelopmental and maturation axis. Along this trajectory, cells in both neuronal clusters (Neuron 1 and 2) demonstrated similar pseudotime maturity scores to previously reported neuronal cultures (Figure 2F).^25^ Collectively, these data indicate that the PSC-derived neurons in this study demonstrated not only a transcriptomic identity consistent with human foetal and PSC-derived MGE-patterned GABAergic neurons, but also a maturity consistent in keeping with previously reported studies.

### Dysregulated transcriptomic landscape in *SGCE*-mutant GABAergic neurons

Differential gene expression analysis was undertaken of the cells in the Neuron1 and Neuron2 clusters to determine potential pathways impacted by *SGCE* mutation. Here, 222 and 112 genes met the threshold (adjusted p value<0.05 and log2 fold change>=0.25) as being significantly up- and down-regulated, respectively, in the *SGCE* -mutation positive neurons compared to their wild-type counterparts (Figure 3A, Supplementary Data 2 and 3). GO term enrichment differed between these two groups of genes, with genes downregulated in mutant neurons enriched for GO terms related to action potential, ion transport, neuronal structure, synaptic signalling and organisation, while genes upregulated in mutant neurons were enriched for terms related to tissue development, cell migration, and response to stimuli (Figure 3B, Supplementary Data 4).

**Figure 3.**
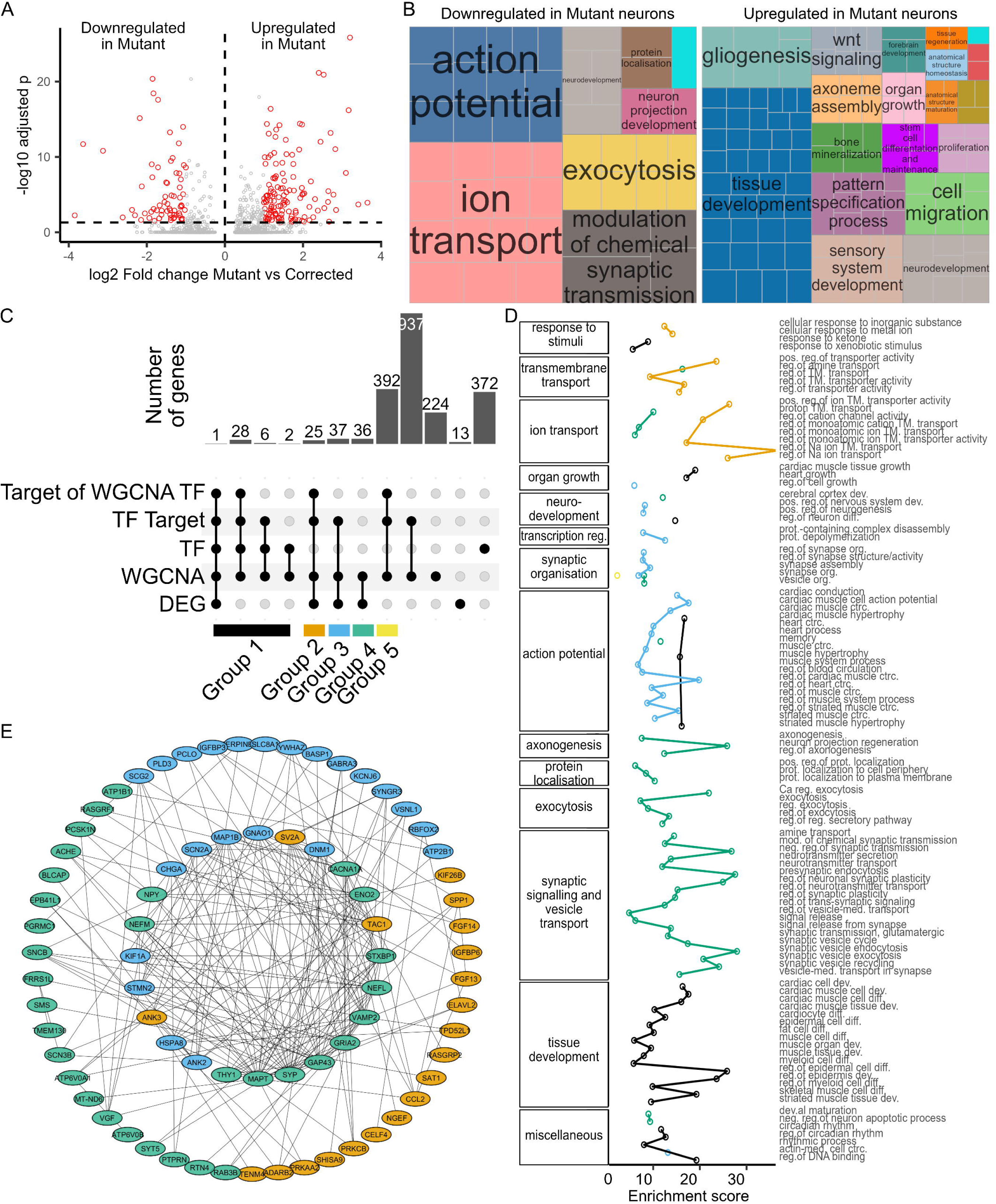
Disrupted transcriptomic landscape of *SGCE*-mutation positive MGE-derived GABAergic inhibitory neurons. **A:** Volcano plot indicating the fold change and adjusted p-values of differentially expressed genes. **B:** Representative biological process gene ontology (GO) terms significantly overrepresented in genes differentially up- and down-regulated in *SGCE* -mutation positive neurons. Similar GO terms are grouped by colour, based on semantic similarity and manual annotation. **C:** Upset plot demonstrating the size of different gene groups categorised according to membership of differentially expressed genes, differentially enriched weighted gene co-expression network analysis (WGCNA) gene modules, and SCENIC regulon analysis. **D:** Biological process GO overrepresentation in five candidate gene groups (Group 1-5). Similar GO terms are grouped together based on semantic similarity and manual annotation. Only GO terms significantly overrepresented are shown (FDR-adjusted p-value>0.05). **E:** Protein-protein interaction network of genes in Group 2-4. Key: Ca: calcium-ion; ctrc.: contraction; DEG: differentially expressed genes; dev.: development; diff.: differentiation; med.: mediated; mod.: modulation; Mut: mutant; Na: sodium; neg.: negative; NPC: neural progenitors; org.: organization; pos.: positive; prot.: protein; reg.: regulated; reg.of: regulation of; RGL: radial glia; TF: transcription factors; TM.: transmembrane; WGCNA: weighted gene co-expression network analysis; WT: wild-type

To identify co-expressed genes correlated with the molecular mechanisms potentially underlying the transcriptomic changes observed in *SGCE* -mutation positive neurons, weighted gene co-expression network analysis (WGCNA) was applied to determine associated gene modules. In total, 33 gene modules were identified, with modules M3, M10, M12, M13, and M18 significantly negatively associated with and differentially enriched in the mutant genotype, while modules M16 and M25 were significantly positively associated with and differentially enriched in the mutant genotype (Supplementary Figure 4A-B, Supplementary Data 5). Interestingly, almost 90% of the 112 DEGs downregulated in mutant neurons form part of the negatively associated and downregulated modules of mutant neurons, whereas <10% of the 222 upregulated DEGs were present in upregulated modules (Supplementary Figure 4C), supporting the view that *SGCE* mutations lead primarily lead to transcriptional downregulation and loss of function.

To gain further insights into the changes underlying the downregulated genes in the *SGCE* -mutation positive neurons, data derived from the DEG, WGCNA and SCENIC regulon enrichment analyses were integrated. Here, 2073 genes were initially shortlisted from the union of i) WGCNA gene modules M3, M10, M12, M13, and M18, ii) transcription factors of SCENIC regulons for which WGCNA gene modules were enriched, and iii) DEGs downregulated in mutant neurons (Figure 2C). Among these genes, five key groups emerged including transcription factors found in differentially enriched WGCNA modules (Group 1), differentially expressed WGCNA genes in regulons of Group 1 transcription factors (Group 2), differentially expressed WGCNA genes in other regulons (Group 3), differentially expressed WGCNA genes not in regulons (Group 4), and other WGCNA genes in regulons of Group 1 transcription factors (Group 5) (Figure 2C, Supplementary Data 6). WGCNA analysis and protein-protein interaction analysis of these gene groups found that Groups 2-4 had the highest mean co-expression weight with genes within and across gene groups, and higher centrality measures in the protein-protein interaction network, but significantly fewer mean number of intragroup co-expression interactions than other groups (Supplementary Figure 5A-B), suggesting that these gene groups are highly associated with the expression level of other genes and may be central to the transcriptomic landscape. GO enrichment analysis revealed that Groups 2-4 were significantly enriched for genes relating to distinct sets of GO terms, namely ion and transmembrane transport (Group 2), neurodevelopment, action potentials and synaptic organisation (Group 3), and axongenesis, protein localisation, exocytosis, and synaptic signalling and vesicle transport (Group 4). By contrast, Group 1 (all transcription factors) was predominantly enriched for GO terms related to action potential and tissue development (Figure 2D; Supplementary Data 7).

These results suggest that *SGCE* mutations disrupt three facets of neuronal function: action potential generation, synaptic and vesicular transport, and axongenesis. To further elucidate the relationship among Group 2-4 genes, a protein-protein interaction network was constructed, based on identification of “hub” genes with the highest measures of centrality measures in each group (Figure 2E, Supplementary Figure 5C; Supplementary Data 8). Group 4 genes demonstrated higher inter-group centrality measures (Supplementary Figure 5C; Supplementary Data 8), suggesting that these genes (enriched for genes related to axongenesis, protein localisation, exocytosis, synaptic signalling and vesicle transport) were likely to functionally link Group 2 and Group 3 genes. “Hub” genes for inter-group interaction in Group 2-4 predominantly involved aspects of axonal organisation and vesicular transport, for example *GNAO1*, *MAP1B*, *STMN2*, *MAPT* (cytoskeletal structure and assembly), *DNM1*, *KIF1A* (transport machinery), *SV2A*, *SYP*, *VAMP2, STXBP1* (synaptic vesicle release), *TAC1*, *CHGA*, *NPY* (encoding precursors of vesicular transport), and *SCN2A*, *GRIA2*, *CACNA1A* (ion channels and receptors for membrane trafficking) (Figure 2E). Interestingly, hub genes from Groups 2 and 3 also included *ANK2* and *ANK3* respectively, and encoding the protein ankyrin, central to the membrane localisation and targeting of ion channels and neurotransmitter receptors.

### Disrupted neurite growth and arborisation in *SGCE*-mutant GABAergic neurons

scRNA-seq analysis implicated that *SGCE* mutations may lead to disrupted axonal organisation and development, potentially manifested in the form of changes to dendritic architecture. Neurite length and branching complexity were examined in day 80 neurons following sparse transfection with a constitutive EGFP expression plasmid (Figures 4A-B), and analysed using the Strahler ordering system and Sholl analysis (Figures 4C-D). Branching pattern analysis identified significantly shorter 1^st^ order branches (Figure 4E; PT1: p=3.30×10^-4^, PT2: p=0.0477, iCas9: p=0.798) but no significant differences in the length of higher order branches (2^nd^ and 3^rd^) in SGCE-mutation positive neurons, compared to their isogenic wild-type controls (PT1: p=3.30×10^-4^, PT2: p=0.0477, iCas9: p=0.798) (Figure 4E). In addition, fewer higher order branches (2^nd^, 3^rd^, 4^th^) were observed across the mutation positive neurons, compared to controls (PT1: p=0.0362, PT2: p=1.20×10^-3^, iCas9: p=0.242) (Figure 4F). Sholl analysis was used to examine the branching complexity of the dendritic architecture, demonstrating that the presence of SGCE mutations was significantly negatively associated with both neurite length and number of branches across all three isogenically matched cell line pairs (PT1 p=1.31×10^-61^, PT2 p=4.11×10^-25^, iCas9 p=9.73×10^-9^) (Figure 4G).

**Figure 4.**
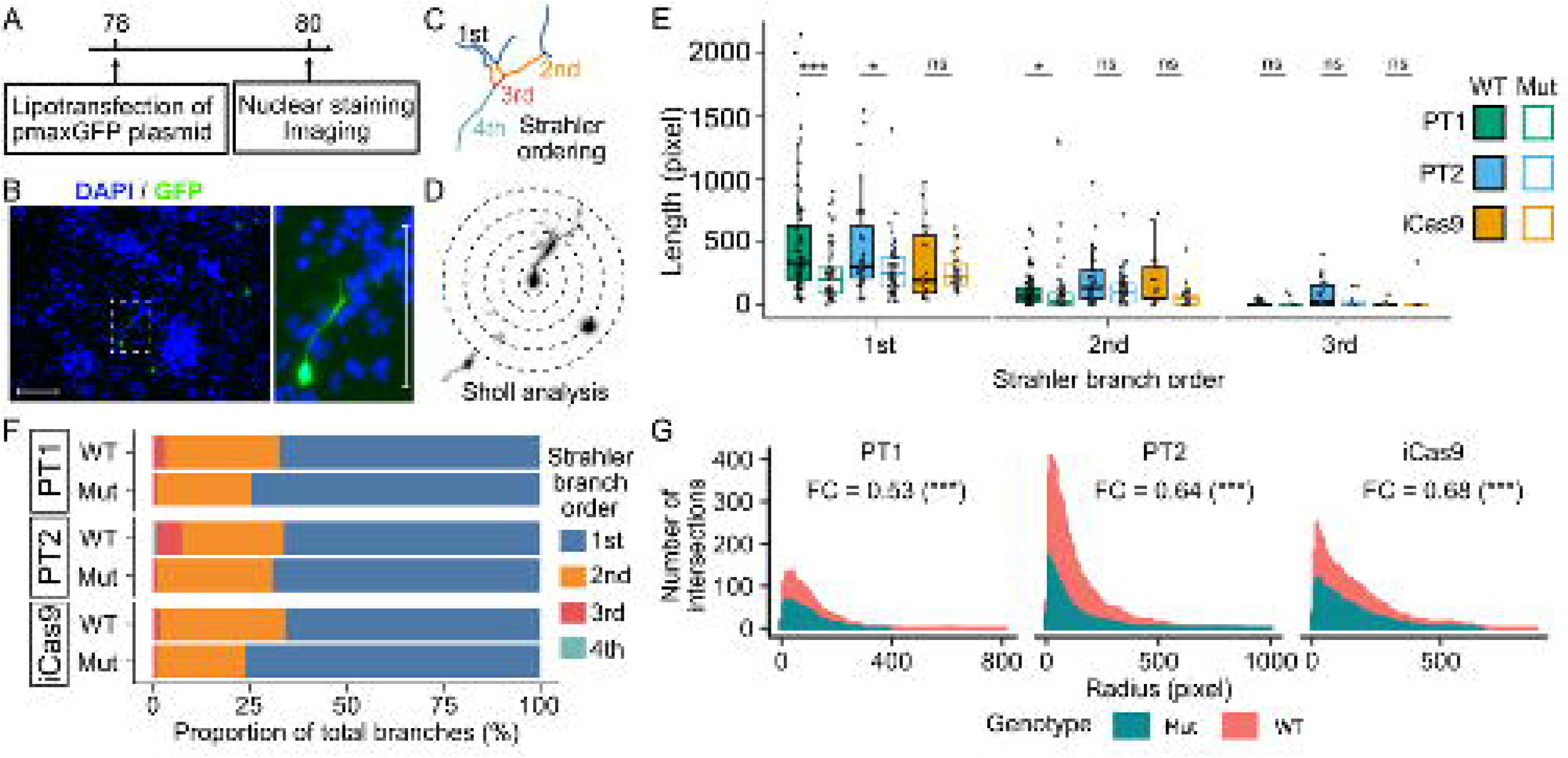
*SGCE* mutations disrupt dendritic architecture with reduced neurite outgrowth and branching. **A-B:** Schematic of the experimental timeline (**A**) and example images (**B**) of neurite tracing. Magnified views for the white box region are shown on the right. Scale bar[=[100[µm. **C-D:** Examples of Strahler branching analysis (**C**) and Sholl analysis (**D**) based on the neuron traced in Panel B. **E:** Box plot representation the median and interquartile values of the length of Strahler order branches. Lines compared using Wilcoxon signed-rank tests with FDR correction. **F:** Horizontal bar plot demonstrating the percentage proportion of each Strahler order branche-type across each cell line. **G:** Sholl analysis histogram of the number of dendritic intersections at sequential radial sizes (steps of 10 pixel size). Fold change (FC) and significance symbols shown are the exponential of the co-efficient and significance for genotype (mutant compared to wild type) of Poisson regression models for each isogenic cell line pair. Key: FC: foldchange; Mut: mutant; PT1: patient #1; PT2: patient #2; WT: wild-type. ns: not statistically significant (p>0.05), *p<0.05. **p<0.01, ***p<0.001.

### Changes to intracellular calcium signalling response observed in *SGCE*-mutant GABAergic neurons

While the changes to ion transport identified during RNA-sequencing analysis may be broad, disruption to calcium signalling has been implicated across multiple monogenic forms of dystonia, including *SGCE*. ^2,26^ As such a high throughput calcium assay, using the FLIPR Penta system, was undertaken on day 80 of differentiation. Compared to control DMSO treatment conditions, overall intracellular peak response calcium concentrations were significantly higher in PSC-derived GABAergic neurons with application of GABA (p<0.001), L-glutamate (p=4.20×10^-4^), and AMPA (p<0.001), but not NMDA (p=0.982) (Figure 4A). Independent co-application GABA and GABA_A_R antagonists bicuculline (p<0.001) and picrotoxin (p<0.001) led to a significant reduction in the GABA-induced calcium response across all cell lines (Figure 5B). The same significant reduction was also observed with co-application of CNQX (AMPA-R antagonist) and L-glutamate (p=0.012), and CNQX aand AMPA (p<0.001). By contrast application of AP5 (NMDA-R antagonist) with L-glutamate identified no significant reduction (p=0.18) (Supplementary Figure 6A). Representative calcium traces are demonstrated in Supplementary Figures 6B and 6C. Significant differences between *SGCE* - mutation positive and wild-type lines were observed only with the GABA-induced response, with significantly larger amplitudes observed in those with wild-type sequence across all three cell line pairs (iCas9 p=0.03, PT1 p=0.02, PT2 p<0.001) (Figure 5B). No significant differences were observed across the cell line pairs in their response to either GABA_A_R antagonists (Figure 5B), potentially indicating that while the number of GABA_A_ receptors are similar between *SGCE* mutant and wild-type genotypes, differences may exist in their secondary response to GABA neurotransmission.

**Figure 5:**
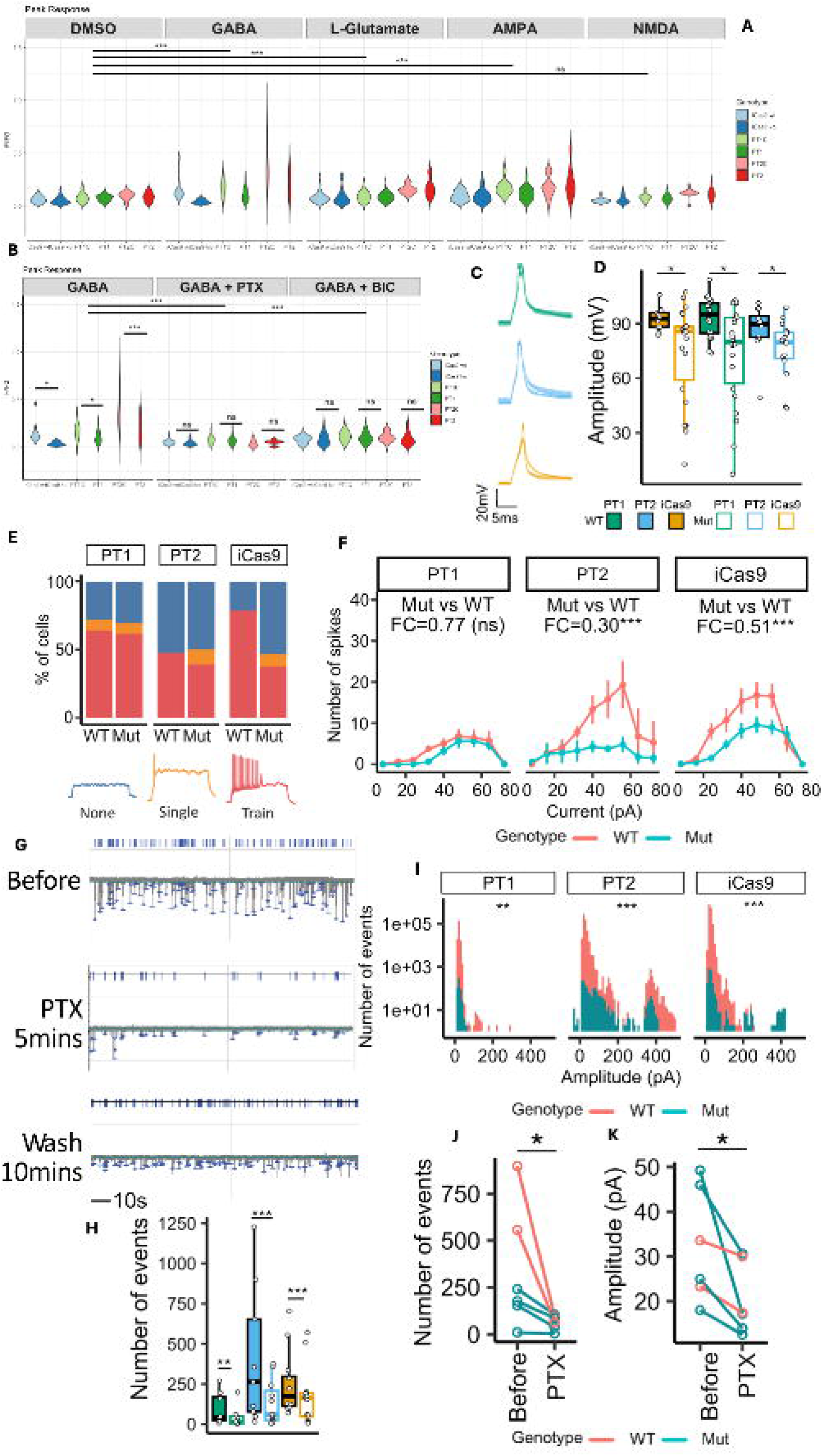
Lower activity and excitability observed in *SGCE* mutation-positive MGE-derived GABAergic neurons compared to wild-type controls. **A-B:** Violin plot comparing the peak calcium fluorescence response (ΔF/F_0_) following small molecule application. Lines compared using one-way ANOVA. **C** Average action potential trace evoked by 5 ms of 200 pA current stimulation. Shaded area represents the standard error of means of all cells. **D:** Box plot representing median and interquartile range of the peak evoked amplitude. Lines compared using Wilcoxon signed-rank tests with FDR correction. **E:** Proportion of cells demonstrating no action potential (blue), single spikes (yellow) and train of spikes (red) following stimulation. **F:** Average number of elicited action potential spikes. Error bars represent the standard error of means of all cells. Fold change (FC) and significance symbols represent the exponential of the co-efficient and significance for genotype (mutant compared to wild-type) in zero-Inflated Poisson regression models of each isogenic cell line pair. **G:** Example trace of spontaneous postsynaptic currents when cells were held at −60 mV at baseline (prior), following PTX application, and following a subsequent wash. **H-I:** The number (H) and amplitude (I) of spontaneous postsynaptic current (SPC) events over 3-minute recordings. Two-sample Kolmogorov-Smirnov test was used to compare the distribution of SPC amplitudes between mutant and wild-type neurons. **J-K** Number (J) and median amplitude (K) of SPC events over 3-minute recordings before and during picrotoxin (PTX) application. Pre- and post-treatment compared using Wilcoxon signed-rank tests with FDR correction. Key: BIC: bicuculline; DMSO: dimethyl sulfoxide; FC: fold change; GABA: γ-aminobutyric acid; Mut: mutant; PT1: patient #1; PT2: patient #2; PTX: picrotoxin; WT: wild-type, *p<0.05. **p<0.01, ***p<0.001. ***: p<0.001; **: p<0.01; *: p<0.05; ns: not significant.

### Reduced activity and excitability in *SGCE*-mutation positive GABAergic neurons

Transcriptomic analysis additionally suggested *SGCE* -mutations to be associated with disruption of action potential generation and synaptic transmission. To further explore potential changes, whole-cell patch clamp analysis was initially undertaken across current- and voltage-clamp modes for all cell lines. No significant differences in passive membrane properties were observed between each *SGCE* mutation carrying line and their isogenic control (Supplementary Figure 7). However, in response to short pulse electric stimulation, action potentials generated from *SGCE* -mutation carrying neurons were of significantly higher amplitude than their WT counterparts (PT1: p=0.0225, PT2: p=0.0281, iCas9: p=0.0281) (Figures 5C-D), while no significant differences were observed across the other measured kinetic metrics (Supplementary Figure 7). With sustained electric stimulation, fewer *SGCE* mutation-positive neurons exhibited a repetitive firing pattern compared to their isogenic control (Figure 5E), while significantly fewer action potential spikes were observed with the mutant genotype following sustained electric stimulation (PT1: p=0.253, PT2: p=2.80×10^-6^, iCas9: p=2.55×10^-6^) (Figure 5F). In addition, *SGCE* -mutation positive neurons exhibited significantly fewer spontaneous postsynaptic currents and lower amplitude than their isogenic matched wild-type controls (PT1: p=9.44 ×10^-3^, PT2: p=1.08×10^-101^, iCas9: p=3.26×10^-68^) (Figure 5G-I), with these spontaneous postsynaptic events likely dependent on GABA_A_R signalling given the reduction in both frequency and amplitude with the application of the GABA_A_R antagonist, picrotoxin (Figures 5G and 5J-K).

Use of multielectrode arrays (MEAs) to evaluate network level neuronal activity, demonstrated an overall increase in activity over time, across all cell lines, indicating progressive development of neuronal functional maturity (Figures 6A-B). The MEA-detected activity was reduced across both mutant and wild-type lines with application of voltage-gated sodium channel blocker, tetrodotoxin (TTX), supporting the biological relevance of the MEA-detected activity given its dependence on voltage-gated sodium channels, a feature of neuroectodermally derived differentiated neurons (Figure 6C).^27^ *SGCE* -mutation positive neurons again demonstrated lower levels of activity with significantly lower spike (PT1: p=1.82×10^-6^, PT2: p=8.64×10^-11^, iCas9: p=4.62×10^-20^) (Figure 6D) and burst frequency (PT1: p=2.57×10^-4^, PT2: p=4.55×10^-13^, iCas9: p=4.73×10^-9^) (Figure 6E), compared to their isogenic wild-type controls. Collectively, *SGCE* -mutation positive GABAergic neurons were less active and excitable, demonstrated fewer spontaneous postsynaptic currents, and lower spike and burst network activity, compared to each of their wild-type controls.

**Figure 6:**
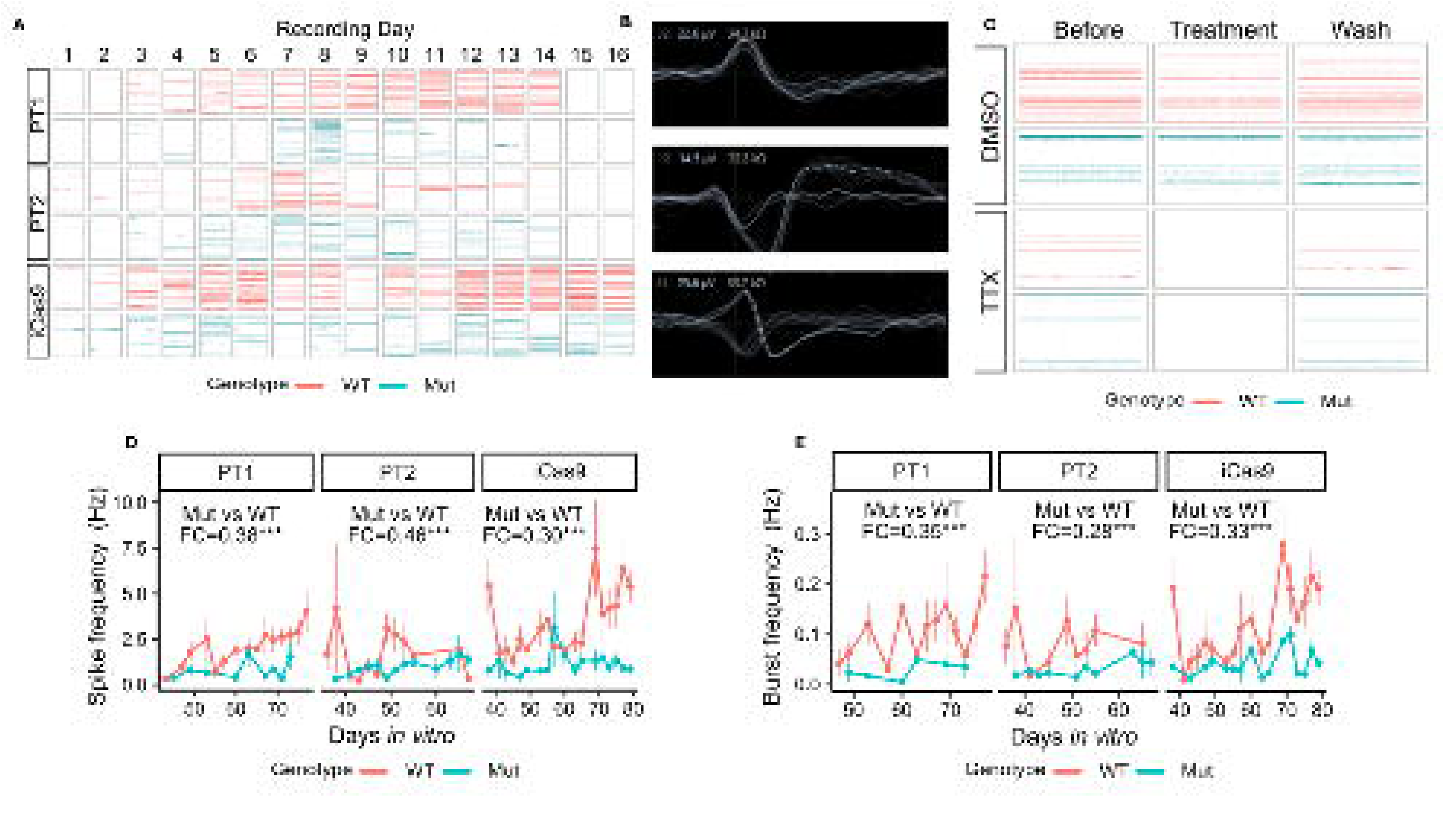
**A-B:** Raster plot (A) and representative waveforms (B) of neuronal activity captured using multi-electrode arrays (MEAs). **C:** Raster plot of neuronal activity before, during tetrodotoxin treatment, and following a subsequent wash. **D:** MEA captured spike frequency across all cell line pairs. **E:** MEA captured burst frequency across all cell line pairs. Data presented as line plots, with each point representing the mean value across a minimum of 3-wells per experiment, from 3 independent experiments per line. Fold change (FC) and significance symbol shown are the exponential of the co-efficient and significance for genotype (mutant compared to wild-type) in gamma regression models for each isogenic cell line pair. Key: Mut: mutant, TTX: tetrodotoxin; WT: wild-type, *p<0.05. **p<0.01, ***p<0.001. ***: p<0.001; **: p<0.01; *: p<0.05; ns: not significant.

## Discussion

Here, we have demonstrated successful differentiation of MGE-patterned GABAergic neurons derived from two patient-derived *SGCE* mutation-positive iPSC lines and a single *SGCE* knock-out human ESC line, coupled with their isogenic WT control lines. The generated neurons resemble human foetal MGE-derived GABAergic neurons when compared with previously reported datasets, both in terms of developmental markers and maturity. Subsequent single cell transcriptomic analysis identified disruption to axonal and cytoskeleton organisation, vesicular transport, synaptic signalling, and action potential generation in the *SGCE* -mutation harbouring lines, compared to controls. Further support for these differences were observed across multiple functional assays including reduced neurite outgrowth and branching complexity, lower intracellular calcium response to GABA stimulation, and a reduction in the neuronal network activity and excitability in the *SGCE* -mutant GABAergic neurons, compared to their wild-type controls. These phenotypes are in contrast with the hyperexcitable phenotype observed in *SGCE* -mutant cortical glutamatergic neurons and are consistent with the loss-of-inhibition hypothesis widely reported in dystonia pathogenesis.^28^

Dystonia is widely considered to be a network disorder, caused by disruption to neuronal activity rather than the loss of specific neuronal subtypes. Consistent with the absence of gross structural abnormalities in MRI studies, we identified no significant differences in the expression of key developmental markers for MGE-derived GABAergic neurons between mutant and control cell lines, in keeping with previous *SGCE* -mutation positive stem cell models differentiated towards cortical glutamatergic neurons and striatal medium spiny neurons.^2^ In addition, scRNA-seq analysis the MGE-derived GABAergic neurons derived in this study found them to resemble their human foetal equivalents as well as a similar transcriptomic maturity to PSC-derived MGE-patterned GABAergic neurons from other studies, supporting the authenticity of the model derived.^25^ However, it should be noted that PSC-derived MGE-patterned GABAergic neurons are likely to be a mixed population of cortical GABAergic interneurons (LHX6, MAF, MAFB, and ERBB4), globus pallidus GABAergic projection neurons (LHX6, NKX2.1, and LHX8), and striatal GABAergic interneurons (LHX6 and NKX2.1). Previous studies have indeed implicated a role for both cortical GABAergic interneurons and globus pallidus projection neurons in the pathogenesis of dystonia with transcranial magnetic stimulation reducing intracortical inhibition in the primary motor cortex,^29^ while intraoperative recordings have demonstrated reduced and irregular firing and burst activities,^30^ and excessive synchronised oscillation in the globus pallidus,^31^ while DBS of the globus pallidus internus has proven to be an effective treatment for dystonia.^32^

Detailed scRNA-seq comparison between *SGCE* -mutant and wild-type neurons identified a network of downregulated genes affecting specific aspects of neuronal function, including axonal and cytoskeletal organization, vesicular transport, synaptic signaling, and action potential generation. More specifically, significantly downregulated genes have been linked with a number of these functions including cytoskeletal organization (*MAP1B*, STMN2, *MAPT*), membrane localization (*EPB41LI, TENM4*) and anchoring of ion channels and receptors (*ANK2*, *ANK3*) supporting the previously hypothesized role for ε-sarcoglycan in these processes.^33^ While others have been linked with synaptic function including vesicle release (*VAMP2*, *SV2A*, *SYP*),^34,35^ and ion channel gating and action potential generation (*SCN2A, CACNA1A, SLC8A1).* ^36,37^ In addition, several of these down regulated genes have also been implicated in clinical disorders linked with neurological and neuropsychiatric phenotypes in which disruption to the balance between excitatory and inhibitory neuronal activity has been implicated, inclusive of dystonia, including *GNAO1*, *STXBP1 and KIF1A*. ^38–40^ Furthermore, the discovery of dystonia-related genes as being dysregulated hubs in *SGCE* -mutant GABAergic neurons implicates a potential convergence of the molecular and cellular mechanisms underlying dystonia, which could be further explored using iPSCs of other monogenic forms of dystonia.

The subsequent structural and functional assays undertaken in this study further support multiple findings from the transcriptomic analysis, including disruption to the dendritic architecture with shorter branches, fewer higher order branches and a less complex branching morphology in *SGCE* mutation positive lines, compared to their isogenic controls. Multiple factors are recognised to impact dendritic morphology, including actin cytoskeletal regulators, neuronal activity, membrane receptors and disruption to calcium signalling,^41–43^ with the lower calcium signalling response following application of GABA, coupled with reduced single-cell and network functional activity in *SGCE* -mutation positive neurons, compared to their wild-type counterparts observed in this study, potentially contributing to this change in morphology. However, as outlined previously, these relationships are likely bidirectional with evidence supporting that changes to dendritic complexity have a secondary impact on neuronal activity and firing patterns.^44^

The multiple functional assays undertaken in this study similarly demonstrated that *SGCE* -mutation positive MGE-derived interneurons to be less active with fewer action potential spikes and post-synaptic currents at a single cell level, while network level MEA analysis demonstrated lower spike and burst frequency, compared to their wild-type isogenic controls. Our transcriptomic analyses provide potential indications as to the factors that may be contributing to these changes, including downregulation of genes involved in the generation of voltage-gated sodium channels (*SCN2A*, *SCN3B*), AMPA receptor components (*GRIA2*, *FRRS1L*) and inwardly rectifying potassium channels (*KCNJ6*), avenues for potential exploration in future work. Evidence increasingly indicates that dystonia arises from disruption to neuronal networks, shaped by an imbalance between excitatory and inhibitory signalling centred around a loss of inhibition.^45,46^ Our previous work has demonstrated that patients derived SGCE-mutation carrying iPSC lines differentiated towards an excitatory glutamatergic lineage harbour a hyperexcitable phenotype, while this study implies that, at a cortical level, this is accompanied by a reduction in inhibitory interneuronal activity.^2^ Given that both of these studies have involved two-dimensional monocultures of each individual cell line, future work will require co-culture of both neuronal subtypes in order to determine ongoing existence of these phenotypes in the other cell types company, and whether their effects are additive or one predominates in a co-culture system.^47^

In conclusion, this study demonstrates reduced excitability in MGE-derived GABAergic neurons harbouring *SGCE* mutations, potentially additive to the observed hyperexcitability in *SGCE* -mutant cortical glutamatergic neurons, in contributing to the hyperkinetic phenotype of Myoclonus Dystonia. Underlying the electrophysiological defects, scRNA-seq revealed dysregulated gene networks centered on specific neuronal functions, including cytoskeletal organization, anterograde trafficking and membrane localization of ion channels and neurotransmitter receptors, consistent with the potential biological function of ε-sarcoglycan in cytoskeletal organization and membrane localization.

Future work will involve a direct assessment of how *SGCE* mutations impact the cortico-basal ganglia-thalamo-cortical neuronal network and the molecular mechanisms by which *SGCE* mutations affect network excitability in different neuronal subtypes.

## Acknowledgements

We would like to thank Meng Li for her ongoing discussion throughout the project, and Michal Rokicki and Joanne Morgan for their work involving iCELL8 library preparation and next generation sequencing, respectively. The computational analysis of this work was undertaken with the support of the Supercomputing Wales project, part-funded by the European Regional Development Fund (ERDF) via the Welsh Government.

## Author contributions

K.P. conceived the study. Z.L. and K.P. designed the experiments. Z.L. performed all experiments (except iCELL8 library preparation and next generation sequencing) with assistance from L.A. O.P. generated iPSC lines used in this project. M.C. and M.L. optimised the original GABAergic neuron differentiation. L.A., M.C., M.L., and K.P. contributed to critical discussion throughout the work. Z.L. and K.P. wrote the manuscript. All authors edited and approved the manuscript.

## Data availability

The sequencing data discussed in this publication have been deposited in NCBI’s Gene Expression Omnibus and are accessible through accession number GSE280716. Scripts used for data analyses are available on GitHub accessible via https://github.com/zongze-li-cardiff/sgce_mge. Raw data are available upon reasonable request.

## Materials and Correspondence

Professor Kathryn Peall, Neuroscience and Mental Health Innovation Institute, Hadyn Ellis Building, Maindy Road, Cardiff, UK, CF24 4HQ.

## Methods

### Experimental Model

Development of both the patient-derived iPSC and CRISPR/Cas9 embryonic stem cell lines have been reported elsewhere.^2^ All participants were examined and confirmed to have a clinical phenotype consistent with Myoclonus Dystonia. They were recruited to the Welsh Movement Disorders Research Network, giving signed informed consent for derivation of iPSC lines in line with the Declaration of Helsinki (REC for Wales, IRAS ID: 146495, REC ref: 14/WA/0017). Confirmation of pathogenic *SGCE* variants was provided by NHS diagnostic laboratory genetic testing for all cases.

### Stem cell culture

iPSCs and hESCs were cultured on Cultrex Stem Cell Qualified Reduced Growth Factor Basement Membrane Extract (R&D Systems) and maintained in Essential 8 flex media (Gibco) under standard culture conditions (37°C, 5% CO_2_). The stem cell media was changed on alternate days and cells passaged every 3-4 days when 70-80% confluency was reached. Primary stem cells (PSCs) were mechanically dissociated and re-plated with Gentle Cell Dissociation Reagent (STEMCELL Technologies) for maintenance and cryopreservation or with ReLeSR (STEMCELL Technologies) for neuronal differentiation upon reaching 70%∼80% confluence.

### MGE-derived GABAergic neuron differentiation

MGE-derived GABAergic neurons were differentiated from PSCs as previously reported.^48^ In brief, differentiation was initiated when the culture reached ∼80% confluence within two days of replating. On day 7 of ESC differentiation or day 9 of iPSC differentiation, cells were mechanically dissociated using 0.5 mM EDTA (Sigma Aldrich) and replated at a ratio of 1:1∼1:1.5 by area. On day 21 and 30, cells were dissociated to single cells (Accutase, 8 minutes at 37°C) and plated at a density of 5×10^5^ cells/cm^2^ and 1.25×10^5^ cells/cm^2^ respectively. 10 µM Y27632 (Stratech Scientific) was added to the culture media for the first day following each passage. Neuronal differentiation was promoted by N2B27-RA basal media supplemented with 100 nM LDN193189 (Sigma-Aldrich), 10 µM SB431542 (Tocris), and 2µM XAV939 (Stratech Scientific) from day 0 to day 9, 200 ng/mL recombinant SHH (Peprotech), and 1 µm Purmorphamine (Sigma-Aldrich) from day 10 to day 20, and 10 ng/mL BDNF (Peprotech) after day 25. On day 25, B27 supplement without RA was replaced by B27 supplement with RA (Gibco).

### Immunocytochemistry

Cultured cells were washed with PBS, fixed in cold (4°C) 3.7% paraformaldehyde (PFA) for 20 minutes, re-washed three times with PBS and stored at 4°C until required. To allow for immunocytochemistry of intracellular markers, cells were permeabilised by sequential washes with 33% and 66% methanol at room temperature, 100% methanol at −20°C, returned to PBS via an inverse gradient and then blocked in PBS-T (0.3% Triton-X-100 in PBS) containing 1% BSA and 3% donkey serum at room temperature for 1 – 3 hours. After blocking, cells were incubated with primary antibodies (Supplementary Table 1) in PBS-T, containing 1% bovine serum albumin and 1% donkey serum, at 4°C overnight. The next day, cells were washed with PBS-T as described above and incubated with secondary antibodies (Supplementary Table 1), in darkness, at room temperature for an hour. The cells were then washed with PBS-T and counterstained with DAPI (Molecular Probes) diluted 1:3000 in PBS-T in darkness. Finally, the cells were washed three times with PBS-T and PBS and mounted with a fluorescence mounting medium. Stained cells were imaged using a Leica DMI6000B inverted microscope, with an average of 10 random fields of view for each staining combination at x20 magnification and images processed using LAS X software (Leica). Automated cell quantification was carried out using Cell Profiler 4.2.7.^49^ Data for immunocytochemical quantification was collected from at least three biological replicates, from at least three independent experiments, for each marker.

### Quantitative real-time PCR

Total RNA was extracted using TRI reagent (Invitrogen), treated with DNase I (Invitrogen)and cDNA generated using EvoScript Universal cDNA Master Version 04 (Roche). Quantitative real-time PCR (qPCR) was performed using Takyon™ Low ROX Probe 2X MasterMix dTTP blue (Eurogentec) to quantify genes of interest. Applied Biosystems QuantStudio™ 7 Flex Real-Time PCR System was used for the standardised qPCR programme. The PCR run involved the temperature being raised from room temperature to 50°C at 1.6°C/second, held at 50°C for 2 minutes, increased to 95°C with an increment of 1.6°C/second, and finally held at 95°C for 10 minutes. The PCR stage involved a total of 45 cycles, each 95°C for 15 seconds, decreasing to 60°C by 1.6°C/second, and held at 60°C for 1 minute for fluorescence data collection. For the melt curve, the temperature was increased to 95°C by 1.6°C/second, held at 95°C for 15 seconds, decreased to 60°C by 1.6°C/second, held at 60°C for 1 minute, increased to 95°C by 0.05°C/second with data collection at every increment, and then held at 95°C for 15 seconds. Dissociation curves were recorded to check for amplification specificity. The threshold of fluorescence detection was automatically determined by the software. Relative quantification was determined using the ΔΔ-C_T_ method with QuantStudio Real-Time PCR Software v1.3 (Thermo Fisher Scientific). All data were normalised to the geometric means of CT values of two endogenous control genes, *GAPDH* and *ACTB*. Reactions were performed in triplicate for each cDNA sample, for each of the three differentiations. Primers are listed in **Supplementary Table 2**.

### Single-cell RNA sequencing

Day 80 neuronal cultures were dissociated with Accutase containing 10 units/mL of papain (Sigma-Aldrich) for 8 minutes at 37°C, resuspended in DPBS (Gibco), mixed with 0.5% bovine serum albumin (Sigma Aldrich) and 50 units/mL DNase I (Sigma Aldrich), and passed through a 35 µm mesh cell strainer. Cells were stained with ReadyProbes™ Cell Viability Imaging Kit, Blue/Red (Invitrogen) following manufacturer’s guidance, and then dispensed into the nano-well plates of the ICELL8® cx Single-Cell System (Takara). Wells containing a single nucleus were automatically and manually selected using the ICELL8 cx CellSelect v2.5 Software (Takara). The sequencing library was prepared using the SMART-Seq ICELL8 application kit (Takara) following manufacturer’s instructions. Next-generation sequencing was performed using the NovaSeq 6000 SP Reagent Kit v1.5 (200 cycles) on an SP flow cell on NovaSeq 6000 (Illumina).

### Single-cell RNA sequencing analysis

Raw sequencing data were converted into FASTQ files containing all indices for each chip which were then demultiplexed using the *cogent demux* command and analysed using the *cogent analyze* command with default settings, as part of the Cogent NGS Analysis Pipeline software 2.0.0 (Takara). Sequences were aligned to the *Homo sapiens* GRCh38.94 primary assembly. The count matrix of each sample was combined in R 4.4.0.^50^ Downstream analysis was performed in R 4.4.0 using Seurat 5.1.0^51^ unless otherwise stated. Only protein-coding genes with at least 5 total counts and expressed in at least 8 cells (1%) were included, and after gene-level filtering, only cells with >2×10^4^ total gene counts, >5000 genes detected, and <10% mitochondrial gene count, were included for downstream analysis.

Raw counts were normalised using the *NormalizeData* function with a scale factor of 1×10^4^ and scaled with the *ScaleData* function regressing out the percentage of mitochondrial gene count. The top 2000 most highly variable genes, identified using the variance-stabilizing transformation method of the *FindVariableFeatures* function, were used for principal component analysis with the *RunPCA* function. The first 20 principal components were used for downstream integration and dimensional reduction, based on the Jackstraw method^52^ and visual inspection of the elbow plot. Data from distinct cell lines were integrated using the *IntegrateLayers* function which tested four different methods (canonical correlation analysis, Harmony, reciprocal PCA, and fast mutual nearest neighbours correction), and yielded similar results. The Harmony method was selected to facilitate standardised analysis with other publicly available datasets ^53^. After data integration, uniform manifold approximation and projection (UMAP) and unbiased Louvain clustering was performed using *RunUMAP, FindNeighbors* and *FindClusters* functions, using the first 20 components of the harmony dimensional reduction matrix and default settings. Differential gene expression analysis was performed using the *FindMarkers* function with Wilcox ranked sum test method and Bonferroni correction.

Gene ontology enrichment analysis was performed using *enrichGO* function in the clusterProfiler 4.12.0 package with all genes in the filtered dataset used as the background dataset. The *Homo sapiens* GO term database was downloaded using the org.Hs.eg.db 3.19.1 package, and rrvgo 1.16.0 was used to group the representative GO terms based on semantic similarity calculated using the *calculateSimMatrix* function. Overlap between genesets were analysed using the *enricher* function in the clusterProfiler 4.12.0 package. Reference mapping was performed using the *FindTransferAnchors* and *TransferData* functions in Seurat based on the first 30 dimensions of the Harmony loadings from the reference dataset. Reference datasets were downloaded from NCBI’s Gene Expression Omnibus and processed in R 4.4.0 using the same filtering threshold and analysis pipeline as described above. Clusters were re-annotated based on the expression of canonical markers and compared to published annotation where possible.

Gene regulatory networks were inferred based on the filtered non-zero raw count using SCENIC 1.3.1 in R 4.3.0. Transcription factor binding motifs were searched within 500-5000bp up- and downs-stream of the transcription start site in the *Homo* sapiens - hg38 - refseq_r80 - v9 databases downloaded from https://resources.aertslab.org/cistarget/databases/. Regulon enrichment score (AUC) calculated by SCENIC was used to test for differential enrichment of regulons and analysed using the Wilcoxon signed-rank test with Bonferroni correction for multiple comparisons.

Weighted correlation network analysis was performed using the hdWGCNA 0.3.01 package ^54–56^. Metacells were created based on seurat_clusters, genotype, and sample variables. The Neuron1 population was used as the group of interest, a soft power threshold of 7 was determined based on the scale free topology model fit, and different genotypes were harmonised during the module eigengene calculation. Differential module eigengene analysis was performed using the Wilcox ranked sum test with Bonferroni correction realised by the *FindDME* function. Gene networks were visualised and analysed in Cytoscape 3.10.2 and in R 4.4.0 with the igraph 2.0.3 package ^57,58^. Protein-protein interaction network data were downloaded from BioGRID v4.4.236, IntAct v247, String v12.0, Reactome release 89, and BioPlex v3, (all accessed on 18 August 2024) and integrated by keeping only unique interaction pairs.^59–64^

### FLIPR calcium assay

Day 30 neurons were plated and cultured according to the standard protocol (Figure 1A). On the day of recording, equal volume of FLIPR Calcium 6 Assay buffer was added to cells without replacing or washing and cells were incubated for 2 hours at 37°C. Drug assay buffers were prepared in HBSS with Ca^2+^ and Mg^2+^ (Gibco) with 20 mM HEPES buffer (Gibco) at 5X concentration. After incubation, cells were imaged on FLIPR Penta system (Molecular Devices) at a frequency of 2 Hz for 1 minute prior to injection. 12.5 µL of the drug assay buffers were dispensed at a speed of 12.5µL/second and images captured at a frequency of 2 Hz for a further 5 minutes. Raw fluorescence intensity data were exported and analysed in R 4.4.0 using a custom script. The average baseline fluorescence intensity for each well was calculated by averaging the raw fluorescence intensity measured prior to drug application. The normalised change in fluorescence intensity above the baseline (ΔF/F0) for each well, at a given timepoint, was calculated using the following formula:

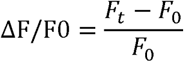

where F_t_ is the fluorescence intensity at timepoint t, F_0_ is the average fluorescence intensity of the baseline period (30 seconds prior to drug application).

### Neurite tracing assay

Day 80 neuronal cultures were sparsely transfected with 80 ng/cm^2^ pmaxGFP™ Vector (Lonza) using Lipofectamine™ 3000 Transfection Reagent (Invitrogen). 0.325 µL/well of Lipofectamine 3000 reagent was used in 25 µL/well final volume for 48-well plates. Following a 24-hour incubation period, the media was fully replaced, and 48 hours after transfection the cells were incubated with NucBlue™ Live ReadyProbes™ Reagent (Hoechst 33342; Invitrogen) in the dark and washed three times with DPBS. Fresh N2B27 basal media, without phenol red, was added for live-cell fluorescence microscopy using the Leica DMI6000B inverted microscope. Neurites were semi-automatically traced using the SNT toolbox in Fiji ImageJ.^65,66^ All neurites of the same cell were anchored to the centroid of the starting point of the primary path. Strahler and Sholl analyses were performed using the SNT toolbox.

### Multi-electrode array assay

Day 30 neurons were plated at a density of 1.25×10^5^ cells/cm^2^ onto CytoView MEA 24 plates (Axion Biosystems) coated with Cultrex, in N2B27 basal media containing 10 ng/mL BDNF and 10 µM Y27632. The day after plating, 1 mL/well of BrainPhys (STEMCELL Technologies), supplemented with 10 ng/mL BDNF, was added to each well and continued throughout the recording period.

Spontaneous extracellular recordings were obtained from 16 electrodes/well using the Axion Maestro Pro system at a sampling rate of 12.5 kHz, 3 kHz Kaiser Window, 200 Hz IIR, and spike detectors. Spikes were detected using the adaptive threshold crossing method with a threshold of 6, pre-spike duration of 0.84 ms, post-spike duration of 2.16 ms, and coincidence event window of 80 µs across 4 electrodes. Cultures were maintained in a 37°C and 5% CO_2_ environment during the recording. Recordings were made three times a week on alternate days from day 36 to day 79 of the differentiation. Activity metrics were calculated based on the list of spikes, bursts, and network bursts, and processed in R 4.4.0 for further analysis. Electrodes with less than 1 spike per minute were omitted.

### Whole-cell patch clamp

Day 30 neurons were plated and cultured on Cultrex coated glass coverslips. Day 78-82 neurons were assayed. Immediately prior to recording the coverslips were transferred to a recording chamber on an Olympus BX61W (Olympus) differential interference contrast (DIC) microscope and continuously perfused at 1.5∼2.5 ml/min with extracellular solution (145 mM NaCl, 10 mM D-glucose, 10 mM HEPES, 3 mM KCl, 2 mM MgCl_2_, and 1.25 mM CaCl_2_, adjusted to pH=7.37∼7.40 with NaOH, and 305∼310 mOsm/L), heated to 35°C. Recordings were performed using a MultiClamp 700B amplifier and 5–7 MΩ resistance pipettes, and filled with intracellular solution (120mM potassium gluconate, 10 mM Phosphocreatine disodium, 10 mM HEPES, 4 mM adenosine 5[-triphosphate disodium, 2 mM MgCl_2_, 2 mM EGTA dipotassium, and 0.5 mM CaCl_2_, adjusted to pH=7.30 with KOH, and 289∼293 mOsm/L). Electrophysiological data were sampled at 20 kHz and filtered at 3 kHz using a Digidata 1550 analogue to digital converter and pClamp 10 software (Molecular Devices). Upon the formation of a GΩ seal, fast and slow components of the pipette capacitance were compensated, and the whole-cell configuration was obtained by applying gentle suction. Resting membrane potential was measured in current clamp mode at 0 pA within the first 20 seconds of whole-cell configuration. Series resistance was monitored throughout the experiment by applying a 10mV pulse at 50 Hz. In current clamp mode, the membrane potential was maintained at −60 mV by injecting a constant current with bridge balance and 4 pF pipette capacitance compensation applied throughout. Single action potentials were evoked by injecting 200 pA current pulse for 5 ms, and repetitive action potential firing evoked by injecting −30 ∼ +70 pA current in 10 pA steps for 1 s.

Spontaneous postsynaptic currents were recorded in voltage clamp mode at −60 mV for an excitatory current or 0 mV for an inhibitory current. Sodium and potassium currents were evoked by a series of −90 ∼ +40 mV in 10 mV steps for 500 ms to determine activation, followed by a 500 ms 40 mV step to assess inactivation, with continuous perfusion of extracellular solution containing 100 nM tetrodotoxin (TTX) or 20 mM tetraethylammonium (TEA), respectively. All recording data, except resting membrane potential data, were analysed in Clampfit 11.2. Resting membrane potential data were analysed in R 4.4.0. Spontaneous postsynaptic current data were filtered with a Bessel (8-pole) lowpass filter, −3 dB cutoff of 2000 Hz and baseline corrected according to the slope between the start and the end of the trace. Peaks were identified using threshold-based event detection with a threshold at three times the standard deviation of the filtered signal, noise rejection of 1 ms, and duration of 1 ∼ 500 ms.

### Statistical analysis

All data were collected from three independent differentiations and presented as mean ± standard error of means unless otherwise specified. All statistical analyses were performed in R 4.4.0. Analyses included one-way ANOVE, Wilcoxon signed-rank test and Kruskal-Wallis test with post-hoc Dunn’s test where appropriate. The distribution of spontaneous postsynaptic current amplitude was analysed using the two-sample Kolmogorov–Smirnov test. False discovery rate (FDR) correction was performed for multiple testing. Sholl analysis data, action potential spike response induced by current squares, MEA spike and burst rate, were analysed with Poisson regression, zero-inflated Poisson regression, and gamma regression models, respectively.

## Supplementary Data

**Supplementary Data 1: Gene modules dynamically regulated along the pseudo-temporal trajectory of integrated pluripotent stem cell-derived datasets.**

**Supplementary Data 2: Differentially expressed genes between** SGCE**-mutation positive and wild-type medial ganglionic eminence-derived GABAergic neurons.**

**Supplementary Data 3: Significantly upregulated and downregulated differentially expressed genes from Supplementary Data 2.**

**Supplementary Data 4. Gene ontology enrichment of differentially expressed genes between*SGCE*-mutation positivej and wild-type medial ganglionic eminence-derived GABAergic neurons.**

**Supplementary Data 5. Genes in weighted gene co-expression network analysis gene modules.**

**Supplementary Data 6. Gene groups based on weighted gene co-expression network analysis, differentially expressed genes, and SCENIC regulon.**

**Supplementary Data 7. Gene ontology enrichment of gene groups.**

**Supplementary Data 8. Centrality measures of protein-protein interaction network of Group 2-4.**

## References

1 Bailey, G. A., Rawlings, A., Torabi, F., Pickrell, O. & Peall, K. J. Adult-onset idiopathic dystonia: A national data-linkage study to determine epidemiological, social deprivation, and mortality characteristics. Eur J Neurol 29, 91–104 (2022). 10.1111/ene.15114

2 Sperandeo, A. et al. Cortical neuronal hyperexcitability and synaptic changes in SGCE mutation-positive myoclonus dystonia. Brain 146, 1523–1541 (2023). 10.1093/brain/awac365

3 Junker, J. et al. Quality of life in isolated dystonia: non-motor manifestations matter. J Neurol Neurosurg Psychiatry (2021). 10.1136/jnnp-2020-325193

4 Albanese, A. et al. Phenomenology and classification of dystonia: a consensus update. Mov Disord 28, 863–873 (2013). 10.1002/mds.25475

5 Balint, B. et al. Dystonia. Nat Rev Dis Primers 4, 25 (2018). 10.1038/s41572-018-0023-6

6 Bailey, G. A., Martin, E. & Peall, K. J. Cognitive and Neuropsychiatric Impairment in Dystonia. Curr Neurol Neurosci Rep 22, 699–708 (2022). 10.1007/s11910-022-01233-3

7 Jinnah, H. A. & Sun, Y. V. Dystonia genes and their biological pathways. Neurobiol Dis 129, 159–168 (2019). 10.1016/j.nbd.2019.05.014

8 Zimprich, A. et al. Mutations in the gene encoding epsilon-sarcoglycan cause myoclonus-dystonia syndrome. Nat Genet 29, 66–69 (2001). 10.1038/ng709

9 Grabowski, M. et al. The epsilon-sarcoglycan gene (SGCE), mutated in myoclonus-dystonia syndrome, is maternally imprinted. Eur J Hum Genet 11, 138–144 (2003). 10.1038/sj.ejhg.5200938

10 Peall, K. J. et al. SGCE and myoclonus dystonia: motor characteristics, diagnostic criteria and clinical predictors of genotype. J Neurol 261, 2296–2304 (2014). 10.1007/s00415-014-7488-3

11 Peall, K. J. et al. SGCE mutations cause psychiatric disorders: clinical and genetic characterization. Brain 136, 294–303 (2013). 10.1093/brain/aws308

12 Esapa, C. T. et al. SGCE missense mutations that cause myoclonus-dystonia syndrome impair epsilon-sarcoglycan trafficking to the plasma membrane: modulation by ubiquitination and torsinA. Hum Mol Genet 16, 327–342 (2007). 10.1093/hmg/ddl472

13 Ettinger, A. J., Feng, G. & Sanes, J. R. epsilon-Sarcoglycan, a broadly expressed homologue of the gene mutated in limb-girdle muscular dystrophy 2D. J Biol Chem 272, 32534–32538 (1997). 10.1074/jbc.272.51.32534

14 Waite, A. J., Carlisle, F. A., Chan, Y. M. & Blake, D. J. Myoclonus dystonia and muscular dystrophy: varepsilon-sarcoglycan is part of the dystrophin-associated protein complex in brain. Mov Disord 31, 1694–1703 (2016). 10.1002/mds.26738

15 Ritz, K. et al. SGCE isoform characterization and expression in human brain: implications for myoclonus-dystonia pathogenesis? Eur J Hum Genet 19, 438–444 (2011). 10.1038/ejhg.2010.206

16 Conte, A. et al. Ten-Year Reflections on the Neurophysiological Abnormalities of Focal Dystonias in Humans. Mov Disord 34, 1616–1628 (2019). 10.1002/mds.27859

17 Mencacci, N. E. et al. Dystonia genes functionally converge in specific neurons and share neurobiology with psychiatric disorders. Brain 143, 2771–2787 (2020). 10.1093/brain/awaa217

18 Tomic, A. et al. Brain Structural Changes in Focal Dystonia-What About Task Specificity? A Multimodal MRI Study. Mov Disord 36, 196–205 (2021). 10.1002/mds.28304

19 Carbon, M. & Eidelberg, D. Abnormal structure-function relationships in hereditary dystonia. Neuroscience 164, 220–229 (2009). 10.1016/j.neuroscience.2008.12.041

20 MacIver, C. L. et al. White Matter Microstructural Changes Using Ultra-Strong Diffusion Gradient MRI in Adult-Onset Idiopathic Focal Cervical Dystonia. Neurology 103, e209695 (2024). 10.1212/WNL.0000000000209695

21 MacIver, C. L. et al. Macro- and micro-structural insights into primary dystonia: a UK Biobank study. J Neurol 271, 1416–1427 (2024). 10.1007/s00415-023-12086-2

22 Martella, G. et al. Impairment of bidirectional synaptic plasticity in the striatum of a mouse model of DYT1 dystonia: role of endogenous acetylcholine. Brain 132, 2336–2349 (2009). 10.1093/brain/awp194

23 Martella, G. et al. Regional specificity of synaptic plasticity deficits in a knock-in mouse model of DYT1 dystonia. Neurobiol Dis 65, 124–132 (2014). 10.1016/j.nbd.2014.01.016

24 Quartarone, A. et al. Abnormal associative plasticity of the human motor cortex in writer’s cramp. Brain 126, 2586–2596 (2003). 10.1093/brain/awg273

25 Allison, T. et al. Defining the nature of human pluripotent stem cell-derived interneurons via single-cell analysis. Stem Cell Reports 16, 2548–2564 (2021). 10.1016/j.stemcr.2021.08.006

26 Charlesworth, G. et al. Mutations in ANO3 cause dominant craniocervical dystonia: ion channel implicated in pathogenesis. Am J Hum Genet 91, 1041–1050 (2012). 10.1016/j.ajhg.2012.10.024

27 Kapucu, F. E., Vinogradov, A., Hyvärinen, T., Ylä-Outinen, L. & Narkilahti, S. Comparative microelectrode array data of the functional development of hPSC-derived and rat neuronal networks. Scientific Data 9, 120 (2022). 10.1038/s41597-022-01242-4

28 Hallett, M. Neurophysiology of dystonia: The role of inhibition. Neurobiol Dis 42, 177–184 (2011). 10.1016/j.nbd.2010.08.025

29 Popa, T. et al. The neurophysiological features of myoclonus-dystonia and differentiation from other dystonias. JAMA Neurol 71, 612–619 (2014). 10.1001/jamaneurol.2014.99

30 Welter, M. L. et al. Pallidal activity in myoclonus dystonia correlates with motor signs. Mov Disord 30, 992–996 (2015). 10.1002/mds.26244

31 Neumann, W. J. et al. A localized pallidal physiomarker in cervical dystonia. Ann Neurol 82, 912–924 (2017). 10.1002/ana.25095

32 Reese, R. & Volkmann, J. Deep Brain Stimulation for the Dystonias: Evidence, Knowledge Gaps, and Practical Considerations. Mov Disord Clin Pract 4, 486–494 (2017). 10.1002/mdc3.12519

33 Menozzi, E. et al. Twenty years on: Myoclonus-dystonia and ε-sarcoglycan — neurodevelopment, channel, and signaling dysfunction. Movement Disorders 34, 1588–1601 (2019). 10.1002/mds.27822

34 Salpietro, V. et al. Mutations in the Neuronal Vesicular SNARE VAMP2 Affect Synaptic Membrane Fusion and Impair Human Neurodevelopment. Am J Hum Genet 104, 721–730 (2019). 10.1016/j.ajhg.2019.02.016

35 Steel, D. & Kurian, M. A. Recent genetic advances in early-onset dystonia. Curr Opin Neurol 33, 500–507 (2020). 10.1097/wco.0000000000000831

36 Zhang, J. et al. Severe deficiency of the voltage-gated sodium channel Na(V)1.2 elevates neuronal excitability in adult mice. Cell Rep 36, 109495 (2021). 10.1016/j.celrep.2021.109495

37 Rinaldi, D. et al. CACNA1A variant associated with generalized dystonia. Neurol Sci 45, 4589–4592 (2024). 10.1007/s10072-024-07592-8

38 Villarroel-Campos, D. & Gonzalez-Billault, C. The MAP1B case: An old MAP that is new again. Developmental Neurobiology 74, 953–971 (2014). 10.1002/dneu.22178

39 Spaull, R. et al. STXBP1 Stop-Loss Mutation Associated with Complex Early Onset Movement Disorder without Epilepsy. Mov Disord Clin Pract 9, 837–840 (2022). 10.1002/mdc3.13509

40 Montenegro-Garreaud, X. et al. Phenotypic expansion in KIF1A-related dominant disorders: A description of novel variants and review of published cases. Hum Mutat 41, 2094–2104 (2020). 10.1002/humu.24118

41 Emoto, K. Signaling mechanisms that coordinate the development and maintenance of dendritic fields. Curr Opin Neurobiol 22, 805–811 (2012). 10.1016/j.conb.2012.04.005

42 Lefebvre, J. L. Molecular mechanisms that mediate dendrite morphogenesis. Curr Top Dev Biol 142, 233–282 (2021). 10.1016/bs.ctdb.2020.12.008

43 Wong, R. O. & Ghosh, A. Activity-dependent regulation of dendritic growth and patterning. Nat Rev Neurosci 3, 803–812 (2002). 10.1038/nrn941

44 Udvary, D. et al. The impact of neuron morphology on cortical network architecture. Cell Rep 39, 110677 (2022). 10.1016/j.celrep.2022.110677

45 Filipovic, S. R. et al. Impairment of cortical inhibition in writer’s cramp as revealed by changes in electromyographic silent period after transcranial magnetic stimulation. Neurosci Lett 222, 167–170 (1997). 10.1016/s0304-3940(97)13370-5

46 Chen, R., Wassermann, E. M., Canos, M. & Hallett, M. Impaired inhibition in writer’s cramp during voluntary muscle activation. Neurology 49, 1054–1059 (1997). 10.1212/wnl.49.4.1054

47 Heider, J. et al. Defined co-cultures of glutamatergic and GABAergic neurons with a mutation in DISC1 reveal aberrant phenotypes in GABAergic neurons. BMC Neurosci 25, 12 (2024). 10.1186/s12868-024-00858-z

48 Cruz-Santos, M., Cardo, L. F. & Li, M. A Novel LHX6 Reporter Cell Line for Tracking Human iPSC-Derived Cortical Interneurons. Cells 11 (2022). 10.3390/cells11050853

49 Stirling, D. R. et al. CellProfiler 4: improvements in speed, utility and usability. BMC Bioinformatics 22, 433 (2021). 10.1186/s12859-021-04344-9

50 R: A Language and Environment for Statistical Computing. v. 4.3.0 (R Foundation for Statistical Computing,, Vienna, Austria, 2023).

51 Stuart, T. et al. Comprehensive Integration of Single-Cell Data. Cell 177, 1888–1902.e1821 (2019). 10.1016/j.cell.2019.05.031

52 Chung, N. C. & Storey, J. D. Statistical significance of variables driving systematic variation in high-dimensional data. Bioinformatics 31, 545–554 (2015). 10.1093/bioinformatics/btu674

53 Korsunsky, I. et al. Fast, sensitive and accurate integration of single-cell data with Harmony. Nature Methods 16, 1289–1296 (2019). 10.1038/s41592-019-0619-0

54 Langfelder, P. & Horvath, S. WGCNA: an R package for weighted correlation network analysis. BMC Bioinformatics 9, 559 (2008). 10.1186/1471-2105-9-559

55 Morabito, S. et al. Single-nucleus chromatin accessibility and transcriptomic characterization of Alzheimer’s disease. Nature Genetics 53, 1143–1155 (2021). 10.1038/s41588-021-00894-z

56 Morabito, S., Reese, F., Rahimzadeh, N., Miyoshi, E. & Swarup, V. hdWGCNA identifies co-expression networks in high-dimensional transcriptomics data. Cell Reports Methods 3 (2023). 10.1016/j.crmeth.2023.100498

57 Csardi, G. & Nepusz, T. The Igraph Software Package for Complex Network Research. InterJournal Complex Systems, 1695 (2005).

58 Shannon, P. et al. Cytoscape: a software environment for integrated models of biomolecular interaction networks. Genome Res 13, 2498–2504 (2003). 10.1101/gr.1239303

59 Chatr-aryamontri, A. et al. MINT: the Molecular INTeraction database. Nucleic Acids Res 35, D572–574 (2007). 10.1093/nar/gkl950

60 Oughtred, R. et al. The BioGRID database: A comprehensive biomedical resource of curated protein, genetic, and chemical interactions. Protein Sci 30, 187–200 (2021). 10.1002/pro.3978

61 del Toro, N. et al. The IntAct database: efficient access to fine-grained molecular interaction data. Nucleic Acids Research 50, D648–D653 (2021). 10.1093/nar/gkab1006

62 Szklarczyk, D. et al. The STRING database in 2021: customizable protein-protein networks, and functional characterization of user-uploaded gene/measurement sets. Nucleic Acids Res 49, D605–d612 (2021). 10.1093/nar/gkaa1074

63 Croft, D. et al. Reactome: a database of reactions, pathways and biological processes. Nucleic Acids Res 39, D691–697 (2011). 10.1093/nar/gkq1018

64 Huttlin, E. L. et al. Dual proteome-scale networks reveal cell-specific remodeling of the human interactome. Cell 184, 3022–3040.e3028 (2021). 10.1016/j.cell.2021.04.011

65 Arshadi, C., Günther, U., Eddison, M., Harrington, K. I. S. & Ferreira, T. A. SNT: a unifying toolbox for quantification of neuronal anatomy. Nature Methods 18, 374–377 (2021). 10.1038/s41592-021-01105-7

66 Schindelin, J. et al. Fiji: an open-source platform for biological-image analysis. Nature Methods 9, 676–682 (2012). 10.1038/nmeth.2019

