## Supplementary material for "Transcriptomic disruption and functional hypoactivity in DYT-*SGCE* MGE-patterned inhibitory neurons"

**Supplementary Figure 1: Canonical marker expression of medial ganglionic eminence-derived GABAergic neuron differentiation.**

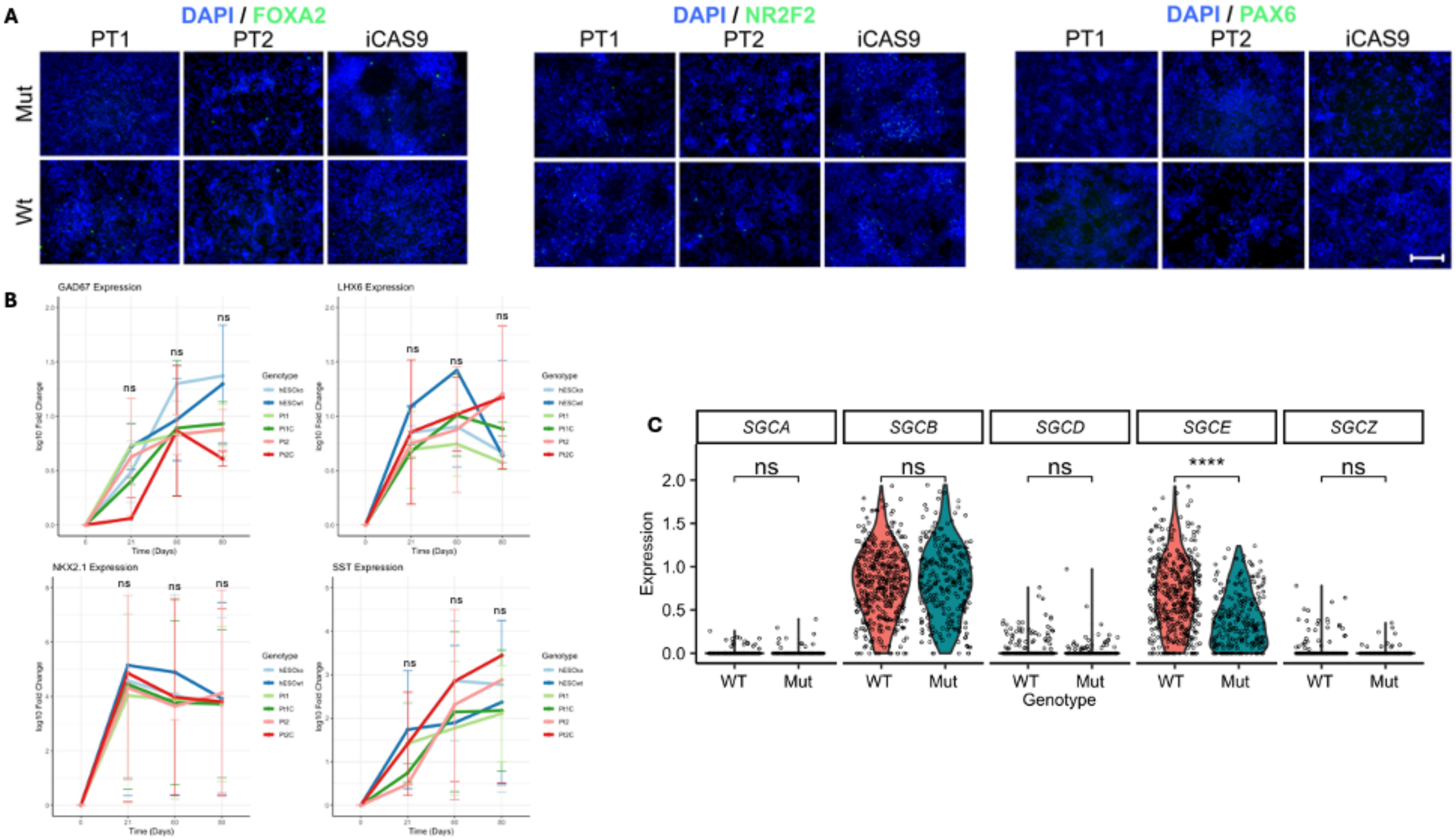


Legend: **A:** Representative immunofluorescence images of the expression of FOXA2 (ventral midbrain marker), NR2F2 (caudal ganglionic eminence marker), and PAX6 (lateral ganglionic eminence and dorsal forebrain marker) on day 25 differentiation. Scale bar = 100 µm. **B:** qRT-PCR expression at Days 0, 21, 60 and 80 for markers of MGE-derived GABAergic neuronal markers, including *GAD67*, *LHX6*, *NKX2.1* and *SST*. Each SGCE mutation carrying line is compared to their wild-type isogenic control. Data presented as mean±SEM from 3 independent experiments per line. Lines compared using two-way ANOVA analysis. **C:** Violin plot showing the normalised expression of genes encodings proteins of the sarcoglycan family in single-cell RNA-sequencing. Lines compared using Wilcoxon signed-rank tests with FDR correction. Mut: mutant; WT: wild-type. *p<0.05. **p<0.01, ***p<0.001, ns: not significant.

**Supplementary Figure 2: Processing and annotation of single-cell RNA sequencing data.**

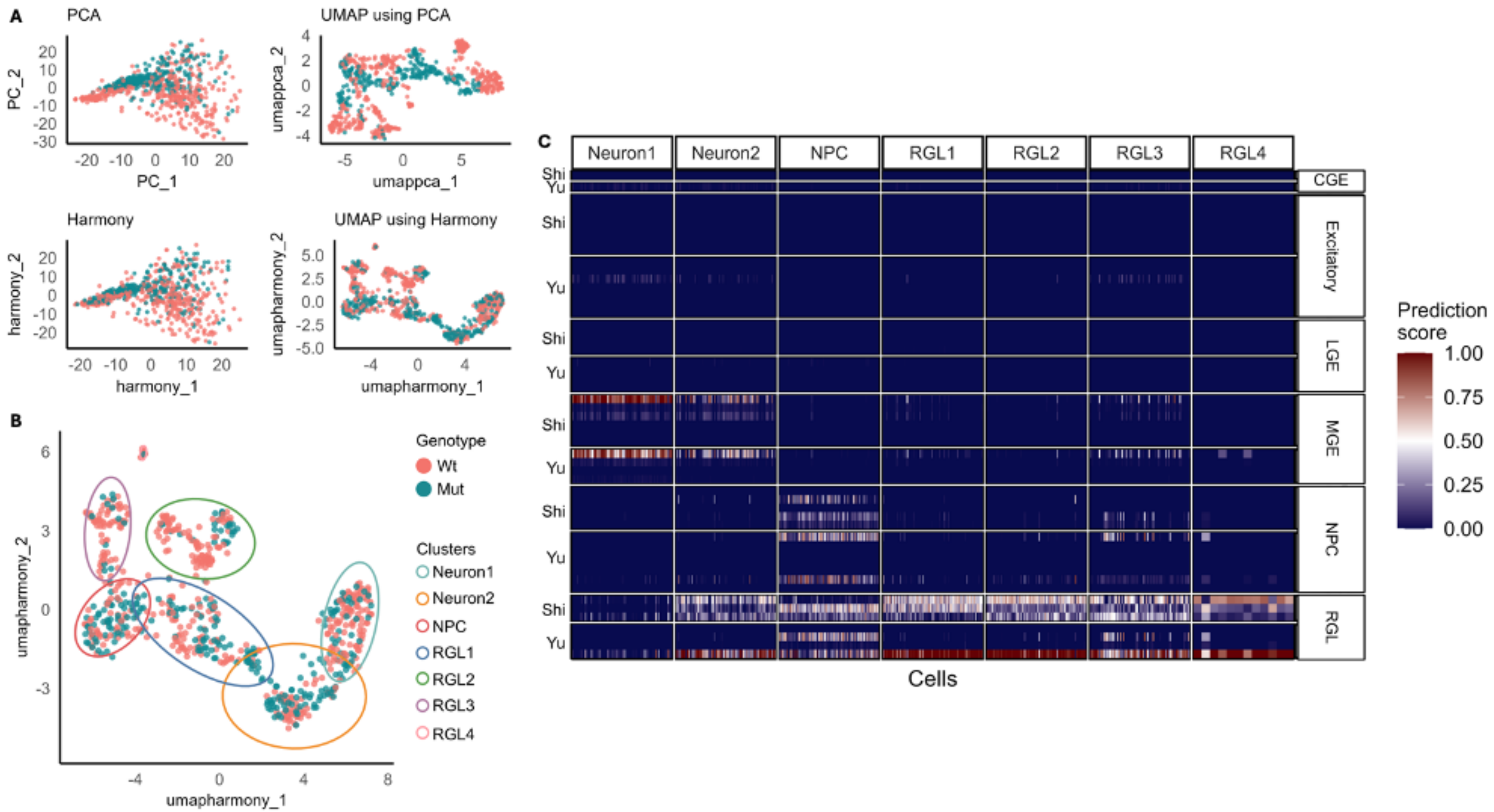


Legend: **A:** Harmony integration aligned wild-type (WT) and mutant (Mutant) cells on Uniform Manifold Approximation and Projection (UMAP) plot coloured by genotype. **B:** D80 UMAP plot of single-cell RNA-sequencing (scRNAseq) of WT and Mut medial ganglionic eminence (MGE)-derived GABAergic neurons. Coloured by genotype and each cluster identified by distinct circumferential colour. **C:** Heatmap of prediction score of reference mapping to human foetal brain datasets (Shi: PMID 34737447; Yu: PMID 34882453). CGE: caudal ganglionic eminence; LGE: lateral ganglionic eminence; MGE: medial ganglionic eminence; Mut: mutant; NPC: neural progenitor; PCA: principal component analysis; RGL: radial glia; UMAP: uniform manifold approximation and projection; WT: wild-type.

**Supplementary Figure 3: Integrated transcriptomic analysis of pluripotent stem cell-derived medial ganglionic eminence-lineage GABAergic neurons.**

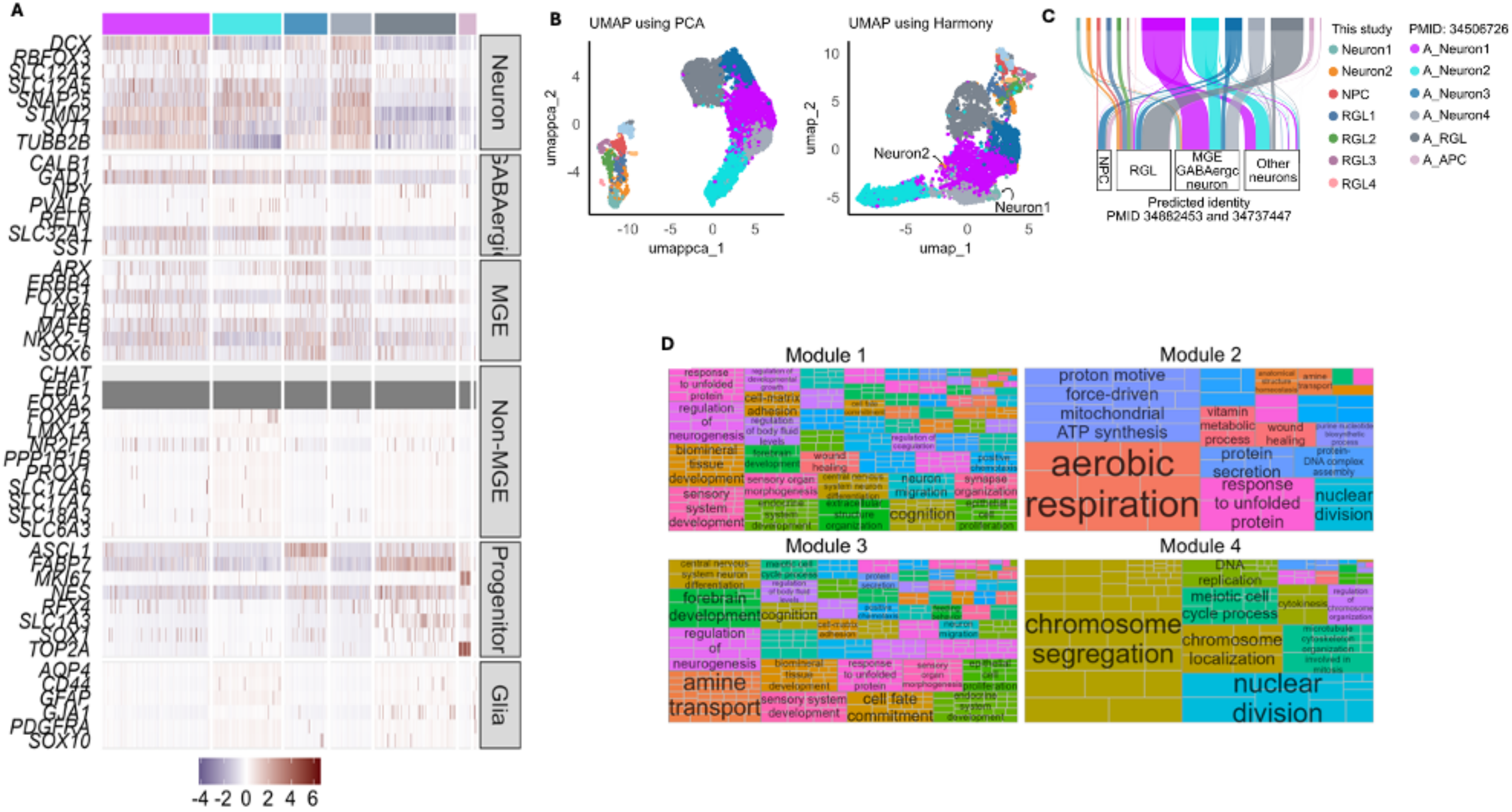


Legend: **A:** Heatmap of the expression of canonical marker genes for distinct lineages and cell types in differing clusters of the Allison *et al.* dataset. **B** UMAP plot before and after Harmony integration of this study and Allison *et al.* dataset (PMID 34506726). **C** Sankey plot demonstrating the predicted identity of the integrated pluripotent stem cell (PSC)-derived dataset based on two published human foetal MGE scRNAseq datasets. **D:** Averaged gene ontology enrichment of gene modules dynamically regulated along the pseudo-temporal trajectory. Similar GO terms are grouped with the same colour based on semantic similarity and manual annotation. **Key:** APC: astrocyte progenitor cell; MGE: medial ganglionic eminence; NPC: neural progenitor; RGL: radial glia; UMAP: uniform manifold approximation and projection.

**Supplementary Figure 4: Single-cell RNA sequencing comparison of wild-type and *SGCE*-mutation positive medial ganglionic eminence-derived GABAergic neurons.**

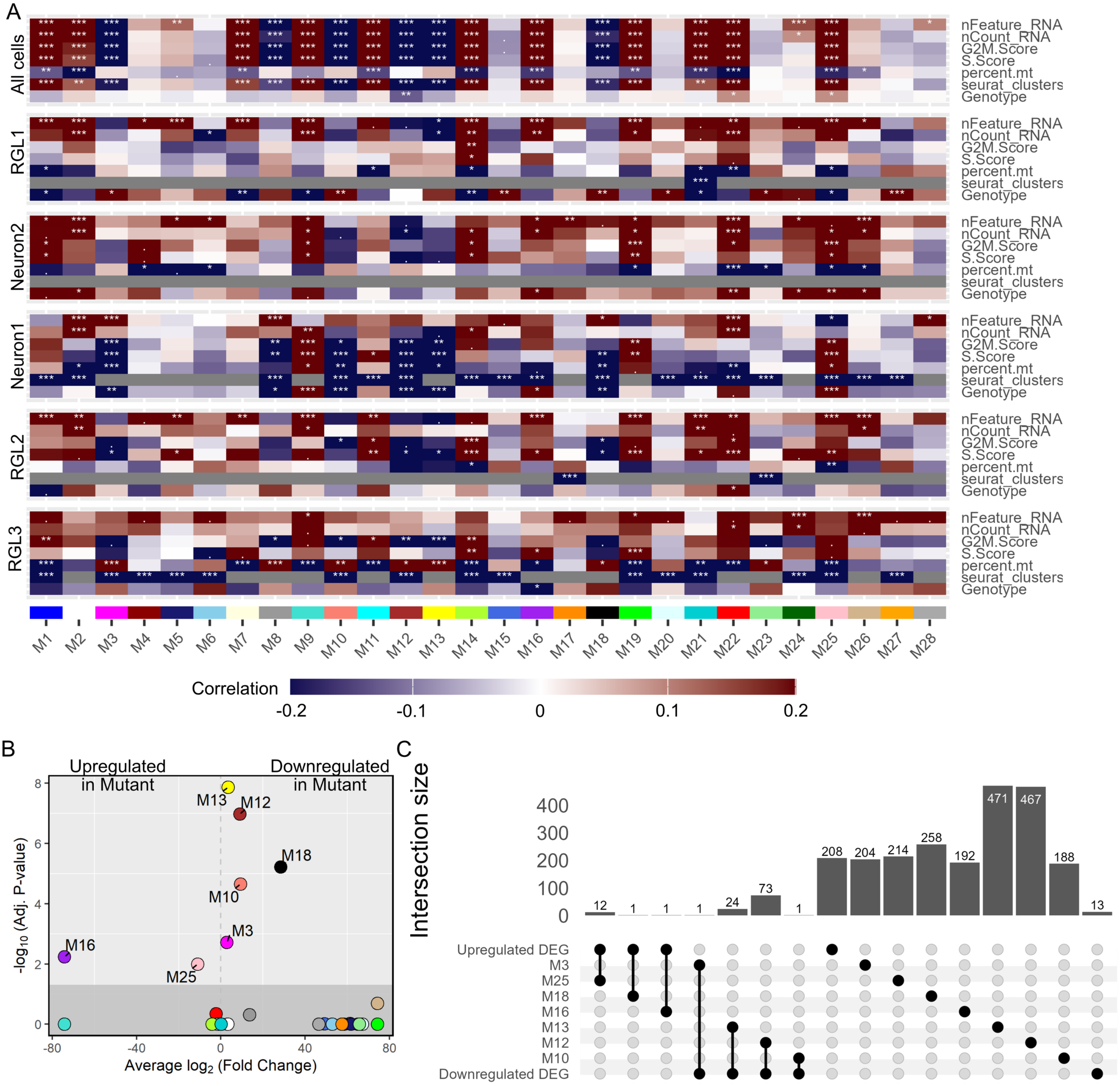


Legend: **A:** Heatmap reflecting the correlation between weighted gene co-expression network analysis (WGCNA) gene modules and different trait variables. **B:** Volcano plot representation of differentially expressed module eigengene analysis, reflecting up- and down-regulated gene modules s individual circles. **C:** Upset plot demonstrating the overlap between differentially expressed gene modules and differentially expressed genes. Key: DEG: differentially expressed genes; Mut: mutant; NPC: neural progenitor; RGL: radial glia; WT: wild type.

Supplementary Figure 5: **Network analysis of candidate gene groups underlying the changes in transcriptomic landscape of *SGCE*-mutation positive neurons compared to wild-type controls.**

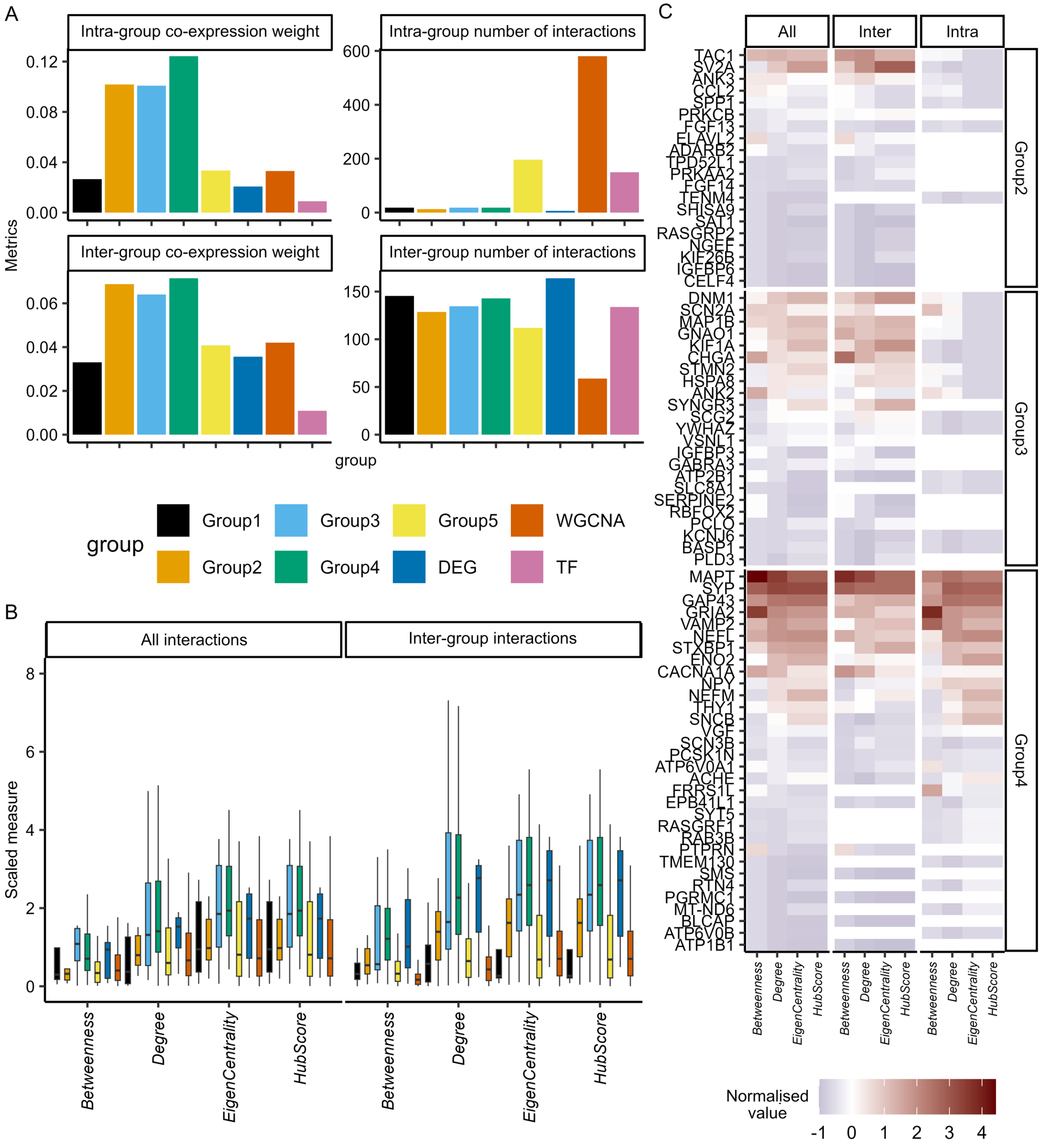


Legend: **A:** Bar plots representing the co-expression weight and number of co-expression edges of the weighted gene co-expression network analysis (WGCNA) across gene groups 1-5, coupled with the DEG, WGCNA and transcription factor groups. **B:** Bar plot comparing network analysis metrics of the protein-protein interaction network across the same distinct gene groups. **C:** Heatmap demonstrating different centrality measures of protein-protein interaction network analysis of Group 2-4 genes. Key: DEG: differentially expressed genes; Inter: inter-group; Intra: intra-group TF: transcription factor; WGCNA: weighted gene co-expression network analysis

**Supplementary Figure 6: Comparison of calcium signalling in wild-type and *SGCE*-mutation positive medial ganglionic eminence-derived GABAergic neurons**.

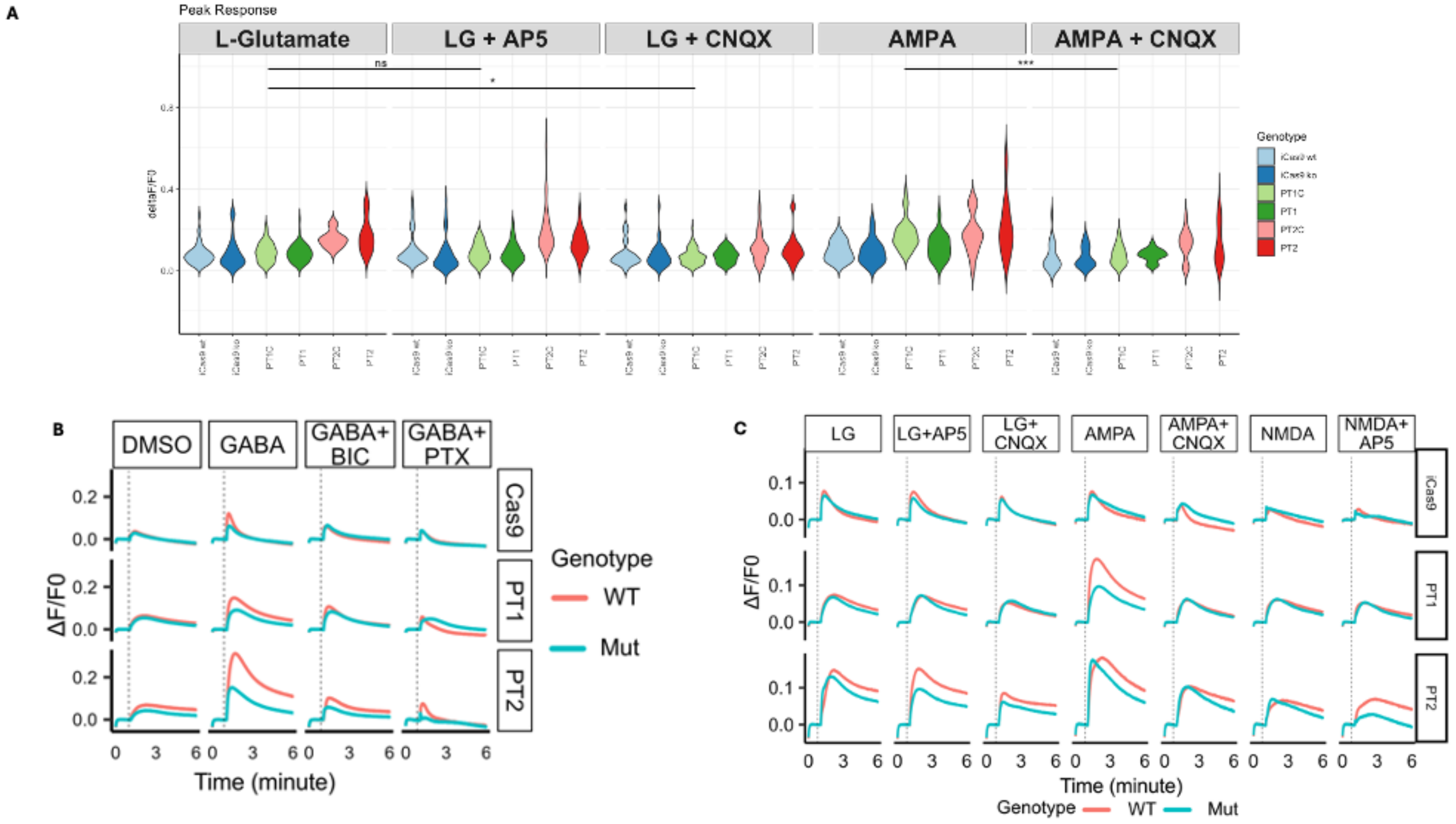


Legend: **A:** Violin plots demonstrating peak calcium fluorescence response ($\Delta F/F_0$) following drug injection. **B and C:** Averaged response trace ($\Delta F/F_0$) derived from the FLIPR calcium assay, with the dashed lines representing drug injection at 60 seconds for DMSO, GABA, GABA + BIC, GABA + PTX (B), and LG, LG + AP5, LG + CNQX, AMPA + CNQX, NMDA, NMDA + APF (C). Key: AMPA: α-amino-3-hydroxy-5-methyl-4-isoxazolepropionic acid; AP5: (2R)-amino-5-phosphonovaleric acid; BIC: bicuculline; CNQX: cyanquixaline; DMSO: dimethylsulfoxide; GABA: γ-aminobutyric acid; LG: L-glutamate; Mut: mutant; NMDA: N-methyl-D-aspartate; PT1: patient #1; PT2: patient #2; PTX: picrotoxin; WT: wild-type Response to small molecule application compared using one-way ANOVA. ***: $p<0.001$; **: $p<0.01$; *: $p<0.05$; ns: not significant

**Supplementary Figure 7: No significant difference in patch-clamp recorded passive membrane properties between SGCE-mutation and wild-type controls.**

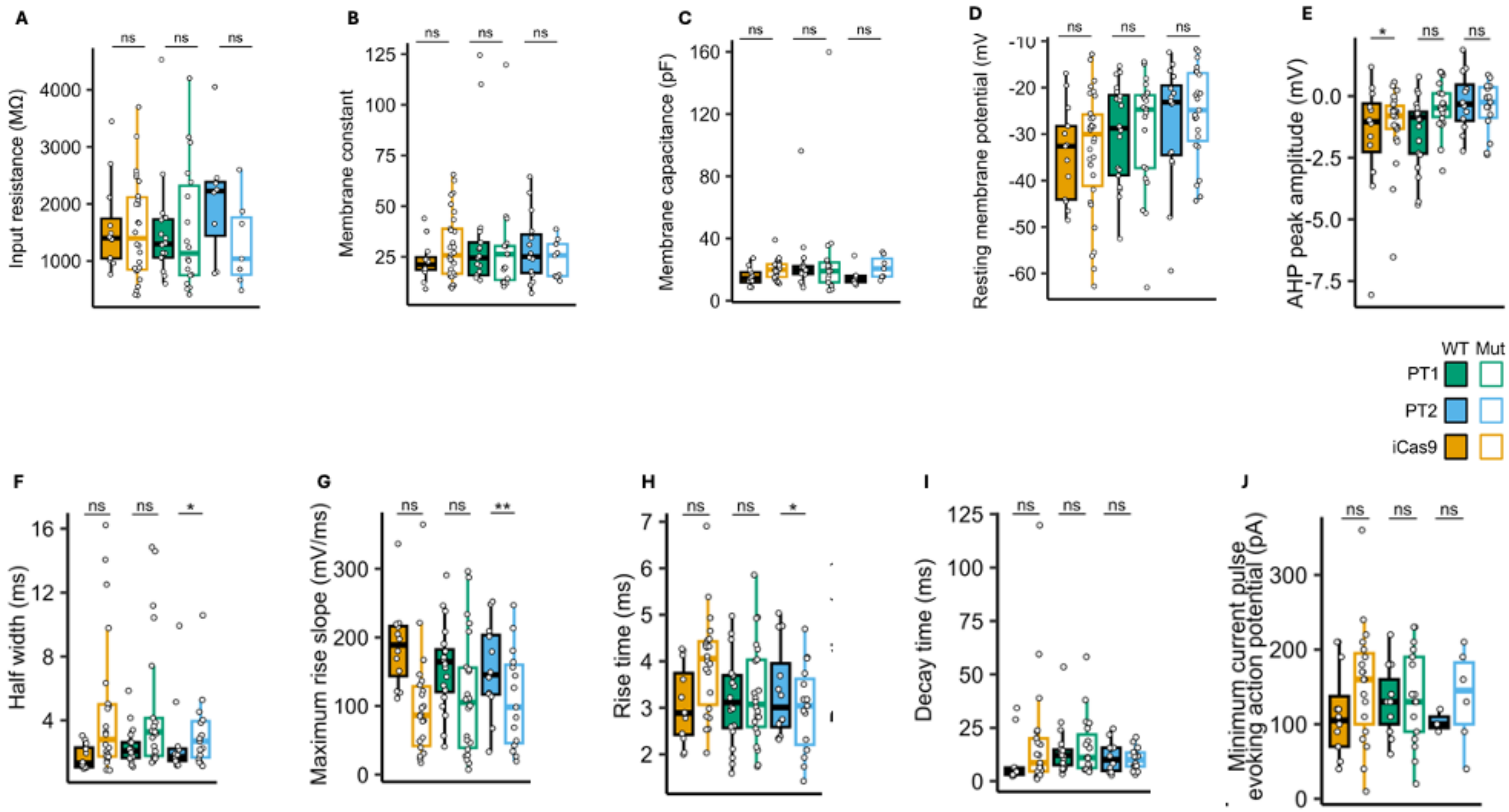


Legend: Box plots representing median and interquartile range values of passive membrane properties, comparing isogenic matched cell line pairs across measures of input resistance (**A**), membrane constant (**B**), membrane capacitance (**C**), resting membrane potential (**D**), AHP peak amplitude (**E**), half-width (**F**), maximum rise slope (**G**), rise time (**H**), decay time (**I**), minimum current pulse evoking action potential (**J**). Key: Mut: mutant; WT: wild-type. Lines compared using Wilcoxon signed-rank tests with FDR correction ***: p<0.001; **: p<0.01; *: p<0.05; ns: not significant.

**Supplementary Tables**

**Supplementary Table 1. Antibodies used in this study.**

| Name | Supplier | Catalogue Number | Dilution used |
|---|---|---|---|
| rabbit anti-CB | Swant | Cat#CB38a | 1:1000 |
| mouse anti-COUP-TFI | Perseus Proteomics | Cat#PP-H8132-00 | 1:1000 |
| mouse anti-COUP-TFII | Perseus Proteomics | Cat#PP-H7147-00 | 1:100 |
| rabbit anti-CR | Swant | Cat#CR7697 | 1:1000 |
| goat anti-FOXA2 | R&D | Cat#AF2400 | 1:1000 |
| rabbit anti-FOXG1 | Abcam | Cat#ab18259 | 1:250 |
| rabbit anti-GABA | Sigma Aldrich | Cat#A-2052 | 1:1000 |
| mouse anti-GAD67 | Millipore | Cat#MAB5406 | 1:500 |
| mouse anti-MAP2 | Sigma Aldrich | Cat#M4403 | 1:500 |
| mouse anti-NESTIN | BD | Cat#BD611659 | 1:1000 |
| rat anti-NEUN-RAT | Abcam | Cat#ab279297 | 1:1000 |
| rabbit anti-NKX2.1 | Abcam | Cat#ab76013 | 1:1000 |
| goat anti-OLIG2 | R&D | Cat#AF2418 | 1:100 |
| rabbit anti-PAX6-RB | Abcam | Cat#AB195045 | 1:1000 |
| rabbit anti-SOX6 | Abcam | Cat#ab30455 | 1:1000 |
| rat anti-SST | Millipore | Cat#MAB354 | 1:50 |
| donkey anti-goat IgG, Alexa Fluor 488 | Invitrogen | Cat#A32814 | 1:1000 |
| donkey anti-goat IgG, Alexa Fluor 555 | Invitrogen | Cat#A32816 | 1:1000 |
| donkey anti-goat IgG, Alexa Fluor 647 | Invitrogen | Cat#A32849 | 1:1000 |
| donkey anti-mouse IgG, Alexa Fluor 488 | Invitrogen | Cat#A21208 | 1:1000 |
| donkey anti-mouse IgG, Alexa Fluor 555 | Invitrogen | Cat#A21202 | 1:1000 |
| donkey anti-mouse IgG, Alexa Fluor 647 | Invitrogen | Cat#A31570 | 1:1000 |
| donkey anti-rabbit IgG, Alexa Fluor 488 | Invitrogen | Cat#A31571 | 1:1000 |
| donkey anti-rabbit IgG, Alexa Fluor 555 | Invitrogen | Cat#A21206 | 1:1000 |
| donkey anti-rabbit IgG, Alexa Fluor 647 | Invitrogen | Cat#A31572 | 1:1000 |
| donkey anti-rat IgG, Alexa Fluor 488 | Invitrogen | Cat#A31573 | 1:1000 |

**Supplementary Table 2. Primers used in this study.**

| Gene | Forward primer | Reverse primer |
|---|---|---|
| GAD67 | CGAGGACTCTGGACAGTAGAGG | GATCTTGAGCCCCAGTTTTCTG |
| GAPDH | ATGACATCAAGAAGGTGGTG | CATACCAGGAAATGAGCTTG |
| LHX6 | ACAGATCTACGCCAGCGACT | CATGGTGTCGTAGTGGATGC |
| NKX2.1 | CGCATCCAATCTCAAGGAAT | TGTGCCCAGAGTGAAGTTTG |
| SST | GCTGCTGTCTGAACCCAAC | CGTTCTCGGGGTGCCATAG |
